# SAX-7/L1CAM acts with the adherens junction proteins MAGI-1, HMR-1/Cadherin, and AFD-1/Afadin to promote glial-mediated dendrite extension

**DOI:** 10.1101/2024.01.11.575259

**Authors:** Elizabeth R. Cebul, Arthur Marivin, Leland R. Wexler, Paola N. Perrat, Claire Y. Bénard, Mikel Garcia-Marcos, Maxwell G. Heiman

## Abstract

Adherens junctions (AJs) are a fundamental organizing structure for multicellular life. Although AJs are studied mainly in epithelia, their core function – stabilizing cell contacts by coupling adhesion molecules to the cytoskeleton – is important in diverse tissues. We find that two *C. elegans* sensory neurons, URX and BAG, require conserved AJ proteins for dendrite morphogenesis. We previously showed that URX and BAG dendrites attach to the embryonic nose via the adhesion molecule SAX-7/L1CAM, acting both in neurons and glia, and then extend by stretch during embryo elongation. Here, we find that a PDZ-binding motif (PB) in the SAX-7 cytoplasmic tail acts with other interaction motifs to promote dendrite extension. Using pull-down assays, we find that the SAX-7 PB binds the multi-PDZ scaffolding protein MAGI-1, which bridges it to the cadherin-catenin complex protein HMP-2/β-catenin. Using cell-specific rescue and depletion, we find that both MAGI-1 and HMR-1/Cadherin act in glia to non-autonomously promote dendrite extension. Double mutant analysis indicates that each protein can act independently of SAX-7, suggesting a multivalent adhesion complex. The SAX-7 PB motif also binds AFD-1/Afadin, loss of which further enhances *sax-7* BAG dendrite defects. As MAGI-1, HMR-1, and AFD-1 are all found in epithelial AJs, we propose that an AJ-like complex in glia promotes dendrite extension.

## INTRODUCTION

A century ago, Warren Lewis wrote, “Were the various types of cells to lose their stickiness for one another […] our bodies would at once disintegrate and flow off into the ground in a mixed stream of ectodermal, muscle, mesenchyme […] and many other types of cells” (Lewis, 1922). This striking visual illustrates the fundamental importance of cell-cell adhesion for multicellular life. Indeed, the acquisition of cell-cell adhesion is thought to be central to the evolution of multicellularity in organisms as diverse as animals, fungi, algae, and bacteria (Niklas, 2014).

The prototypical and best-studied animal cell adhesion complexes are those found in epithelia. Two main adhesion complexes – tight junctions (TJs) and adherens junctions (AJs) – join epithelial cells together into a continuous sheet that separates the inside of an organism from the outside world (Farquhar and Palade, 1963; Hartsock and Nelson, 2008). TJs act as a gate at the interface between the outward-facing ‘apical’ surface of the epithelium and the inward-facing ‘basolateral’ one, blocking diffusion between these compartments. AJs, on the other hand, provide structural integrity to the epithelial sheet by adhering cells to one another (Farquhar and Palade, 1963; Hartsock and Nelson, 2008; Niessen, 2007). Importantly, AJs connect to the actin cytoskeleton, thus creating a unified actin network that spans the epithelium and mechanically couples cells.

The core AJ transmembrane cell adhesion molecules, classical cadherins, are well conserved across phyla (Abedin and King, 2010; Kemler, 1992; Lynch and Hardin, 2009; Takeichi, 1995). To confer cell adhesion, cadherins on neighboring cells interact in *trans* via homophilic binding of their ectodomains. Intracellularly, cadherins connect to the cytoskeleton by binding to β-catenin, which in turn interacts with the actin-binding protein α-catenin and thus forms a link to underlying actin networks (Shapiro and Weis, 2009). This core cadherin-catenin complex associates with a variety of scaffolding proteins that may bolster structural integrity, fortify connections to the cytoskeleton, and facilitate interactions with regulatory signaling pathways (Houssin et al., 2015; Mandai et al., 2013; Niessen and Gottardi, 2008; Stetak et al., 2009).

Interestingly, AJs have also been observed outside epithelial tissues, including in the nervous system. For example, autotypic contacts between paranodal loops of myelinating glia are ultrastructurally similar to epithelial AJs (Fannon et al., 1995). Meanwhile, other neuron-glia attachments appear ultrastructurally distinct from epithelial AJs (Spacek, 1985; Ventura and Harris, 1999), raising the question of whether they utilize a unique set of proteins or if they repurpose epithelial adhesion proteins to build more diverse structures. Pioneering studies of cell adhesion complexes described a remarkable diversity of ultrastructural morphology across tissues, suggesting that adhesion complexes may have the flexibility to take on a range of forms beyond what is observed in model epithelia grown *in vitro* (Farquhar and Palade, 1963). Here, we use the power of *C. elegans* genetics to identify factors that control specialized neuron-glia attachments and find substantial molecular overlap with the classical components of epithelial AJs, suggesting that the core AJ machinery is repurposed to mediate neuron-glia interactions.

*C. elegans* offers several advantages to study neuron-glia attachment: each neuron and glial cell is uniquely identifiable and develops highly stereotyped cell-type-specific attachments, and an impressive toolbox of cell-specific promoters is available that allows single neurons and glia to be visualized and manipulated *in vivo* (Lamkin and Heiman, 2017; Mizeracka and Heiman, 2015; Singhvi and Shaham, 2019; Fung et al., 2020). We focused on specialized attachments between two sensory neurons, called URX and BAG, and a single glial cell called the lateral IL socket (ILso) (Fig. 1A). Most *C. elegans* sensory neurons extend dendrites that protrude through a glial tube to gain direct access to the external environment (Ward et al., 1975). Their dendrites form ring-shaped TJs and AJs with the glia, separating an outward-facing apical region of the dendrite from an inward-facing basolateral region (Lillis et al., 2022; Low et al., 2019). These sensory neurons thus resemble cells in sensory epithelia, such as olfactory neurons in the olfactory epithelium, taste cells in taste buds, and hair cells in the inner ear (Heiman, 2022). By contrast, URX and BAG dendrites are not exposed to the external environment (Ward et al., 1975) and, while they form elaborate membranous attachments to the lateral ILso glia, they are not wrapped by this glial cell and ultrastructurally do not appear to have TJs or AJs (Cebul et al., 2020). Thus, we reasoned that these neuron-glia attachments may more closely model neuron-glia contacts found in the vertebrate central nervous system.

**Figure 1.**
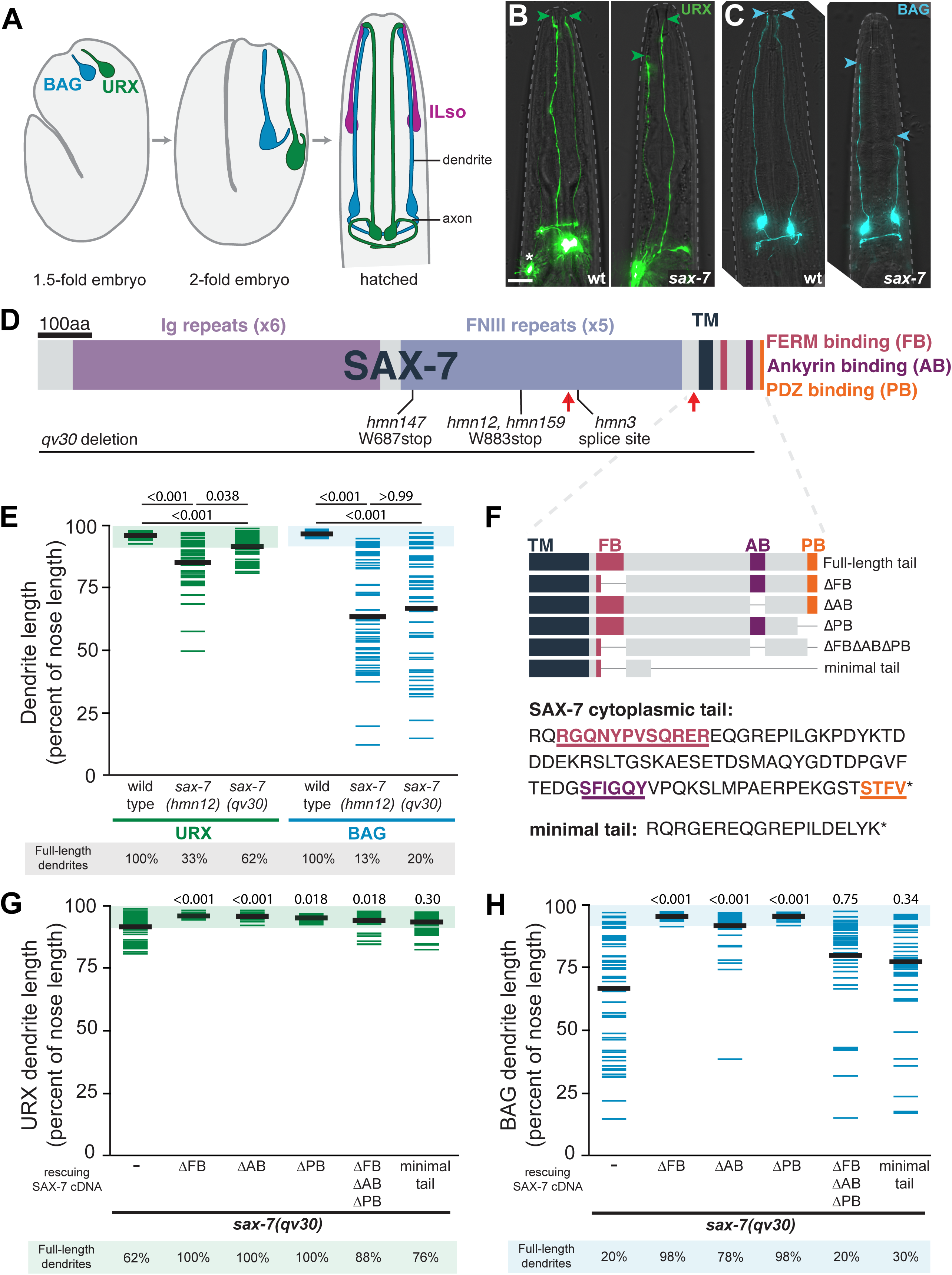
The SAX-7 cytoplasmic tail is required for URX and BAG dendrite extension. (A) Schematic depicting URX (green) and BAG (blue) dendrite extension starting at the 1.5-fold stage (left) and ending at hatching (right). Early in development, the dendrite endings anchor to the presumptive nose, and the dendrites are stretched as the embryo elongates. In the mature structure, URX and BAG dendrite endings attach to the ILso glial cell (magenta). In mutant animals where dendrite endings fail to remain attached to the nose, the dendrites appear truncated and do not extend to the nose tip. For simplicity, only one of each of the bilaterally symmetric URX and BAG neurons are shown in embryo schematics. (B-C) Wild-type (left) and *sax-7(qv30)* (right) L4 larvae expressing cell-specific reporters for URX (B, green, *flp-8*pro) and BAG (C, blue, *flp-17*pro) neurons. Dendrites fail to extend to the nose in *sax-7* mutants. Anterior, up. The dashed line outlines the head. Green and blue arrowheads, URX and BAG dendrite endings, respectively. Asterisk indicates additional cell (AUA) labeled by *flp-8*pro. Scale bar, 10 µm. (D) SAX-7 protein schematic indicating relevant domains, motifs, and alleles. Red arrows indicate putative metalloprotease cleavage sites. TM, transmembrane. (E) URX (green) and BAG (blue) dendrite lengths quantified as a percentage of the distance from the cell body to the nose in the indicated genotypes. Each colored bar represents a single dendrite; black bars indicate population averages. The shaded region marks wild-type mean ± 5 SD, and the percentage of dendrites in this range (“full-length”) is indicated below the plots. *sax-7(hmn12)* data is reproduced from (Cebul et al., 2020). (F) Schematic of the SAX-7 cytoplasmic tail indicating the deletion constructs used in G-H. FB, FERM binding; AB, ankyrin binding; PB, PDZ binding. (G-H) Quantification of URX (G) and BAG (H) dendrite lengths in *sax-7(qv30)* animals bearing no transgene (–) or in animals expressing a SAX-7 cDNA with the indicated deletions under a broad embryonic promoter (*grdn-1*pro). Control (no transgene) data are the same as in (E). p-values, Kruskal-Wallis with Dunn’s post hoc test and adjusted for multiple comparisons. For E, G, and H, n≥46 dendrites per genotype.

We previously showed that URX and BAG dendrites develop by retrograde extension, in which the nascent dendrite endings anchor at the presumptive nose early in development and dendrites are later stretched to their full lengths during embryo elongation (Fig. 1A) (Cebul et al., 2020). If dendrites fail to anchor to the nose, they do not fully extend. Using forward genetics, we found that the neuron-glia cell adhesion molecule SAX-7/L1CAM, as well as the scaffolding protein GRDN-1/CCDC88C, are required for URX and BAG dendrite extension (Cebul et al., 2020). Both factors can act in glia to promote dendrite extension, suggesting that URX and BAG dendrite endings are anchored to the nose during embryogenesis by attachments to glia. Here, we further exploit this system to identify additional factors that promote dendrite-glia attachment, either in parallel or together with GRDN-1 and SAX-7. Through a candidate gene approach, we identify requirements for the scaffolding protein MAGI-1 and the classical cadherin HMR-1, and show that MAGI-1 can physically bridge SAX-7 to the cadherin-catenin complex. We also identify a contributing role for the actin binding protein AFD-1/Afadin, which was previously shown to interact with MAGI-1 and the cadherin-catenin complex in epithelial AJs. Our results lead to the intriguing possibility that ancient epithelial adhesion machinery may have been repurposed by the nervous system to build neuron-glia attachments.

## RESULTS

### The SAX-7 cytoplasmic tail is required for URX and BAG dendrite extension

To better understand the molecules that mediate neuron-glia attachments, we focused on SAX-7 for several reasons: (i) it is a conserved, cell-surface transmembrane neural adhesion molecule; (ii) we had isolated multiple alleles of it in genetic screens for mutants that disrupt URX and BAG dendrite extension; and (iii) our mutants were partially rescued by re-expression of SAX-7 in either neurons or glia and were almost fully rescued by re-expression in both neurons and glia together (Cebul et al., 2020), suggesting that SAX-7 may promote URX and BAG dendrite extension by directly mediating neuron-glia adhesion.

Interestingly, SAX-7 was recently shown to undergo proteolytic cleavage *in vivo* and to function, at least in some contexts, in its cleaved form (Desse et al., 2021). All of our mutant alleles are predicted to produce truncated SAX-7 proteins reminiscent of the cleaved SAX-7 products (Fig. 1D), suggesting they may retain partial activity. We therefore examined a molecular null allele of *sax-7* generated by CRISPR/Cas9-mediated deletion of the entire *sax-7* locus (*sax-7(qv30),* Fig. 1D) (Desse et al., 2021). We found that the molecular null allele *sax-7(qv30*) causes BAG dendrite extension defects with expressivity and penetrance comparable to our truncation allele *sax-7(hmn12*) (Fig. 1C,E; percent of dendrites that are full-length, 20% *qv30*, 13% *hmn12* (Cebul et al., 2020)). In contrast, the URX dendrite extension phenotype is less severe in the null than in truncation alleles (Fig. 1B,E; percent of dendrites that are full-length, 62% *qv30*, 33% *hmn12* (Cebul et al., 2020)). This difference suggests that truncated SAX-7 gene products may interfere with the function of SAX-7 interactors in some contexts.

With a null allele of *sax-7* in hand, we sought to gain insight into how SAX-7 promotes URX and BAG dendrite extension. We reasoned that a likely role for SAX-7 is to provide structural integrity to neuron-glia contacts by coupling extracellular adhesion to the cytoskeleton. In addition to its extracellular adhesion domains, SAX-7 contains a cytoplasmic tail with FERM-binding (FB), ankryin-binding (AB), and PDZ-binding (PB) motifs that can interact with a variety of cytoskeletal-associated molecules (Fig. 1F) (Zhou et al., 2008). To test the importance of these motifs, we introduced transgenes bearing versions of SAX-7 with targeted cytoplasmic tail deletions into the null *sax-7(qv30)* background (Fig. 1F). We then tested how well each transgene rescued URX and BAG dendrite extension defects. We found that deleting each of the known interaction motifs individually had little or no effect on rescue (Fig. 1G,H, ΔFB, ΔAB, ΔPB). By contrast, deleting these binding motifs in combination or using a portion of the SAX-7 cytoplasmic tail that provides only the minimal signals necessary for subcellular trafficking (Lillis et al., 2022) each failed to rescue BAG dendrite extension defects (Fig. 1G,H). Similar trends were observed for URX, although it is more difficult to detect changes in rescuing activity because the *sax-7* phenotype is milder. These results suggest that the SAX-7 cytoplasmic tail may engage with multiple interactors in a partially redundant manner to support dendrite extension, possibly as part of a larger adhesion complex. We therefore turned our attention to possible SAX-7 interactors that may contribute to URX and BAG dendrite extension.

### URX and BAG dendrite extension requires the SAX-7-interacting protein MAGI-1

Relatively little is known about the protein complexes that interact with the SAX-7 cytoplasmic tail, and different partners may interact with SAX-7 in different developmental contexts. In the context of neuronal cell body positioning, SAX-7 genetically interacts with UNC-44/ankryin and STN-2/γ- syntrophin and physically binds them through its cytoplasmic AB and PB motifs, respectively (Zhou et al., 2008). However, we previously found that neither UNC-44 nor STN-2 are required for URX and BAG dendrite extension (Cebul et al., 2020), suggesting they may not be relevant in this context. SAX-7 acts with another partner, MAGI-1, to promote epithelial cell adhesion during embryogenesis (Lynch et al., 2012). To test whether interactions with MAGI-1 are relevant to URX and BAG dendrite extension, we examined two previously described *magi-1* mutants (Fig. 2A). We found that URX and BAG dendrites fail to extend in *magi-1* mutants, similar to defects seen in *sax-7* and *grdn-1* mutants (Fig. 2B,C,F,G).

**Figure 2.**
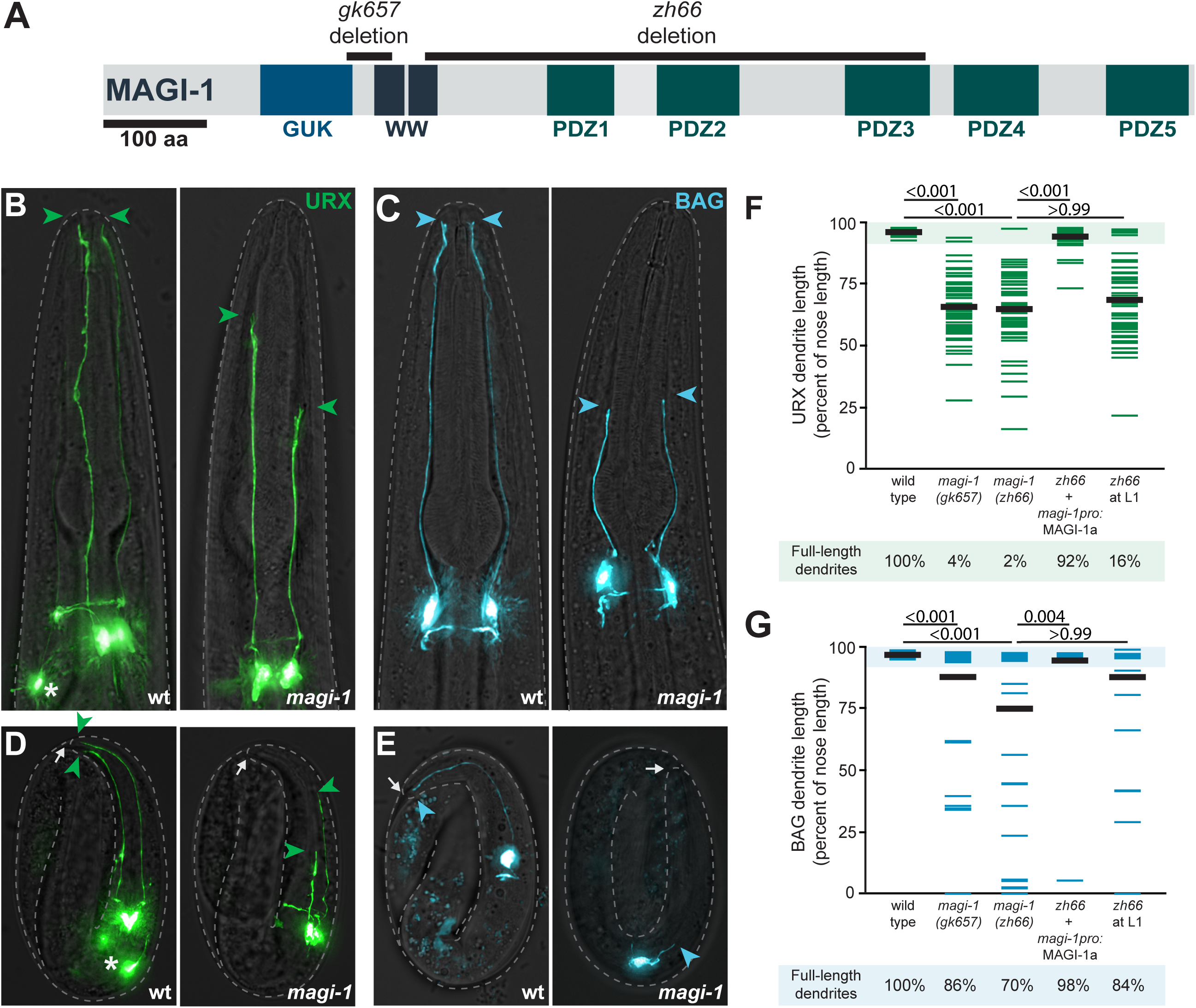
URX and BAG dendrite extension requires MAGI-1. (A) Schematic of MAGI-1 showing conserved domains and alleles used in this study. The guanylate kinase (GUK) domain is likely catalytically inactive. (B-E) URX (B,D, green, *flp-8*pro) and BAG (C,E, blue, *flp-17*pro) neurons in wild-type (left) and *magi-1(zh66)* (right) animals. Green and blue arrowheads, URX and BAG dendrite endings, respectively. Asterisk indicates additional cell (AUA) labeled by *flp-8*pro. (B,C) Images of L4 larvae showing that dendrites fail to extend to the nose in *magi-1* mutants. Anterior, up. The dashed line outlines the head. (D,E) Images of embryos showing that URX (D) and BAG (E) dendrite extension phenotypes are present before hatching. Approximate outline of each embryo is indicated by the dashed line, and the nose is marked with a white arrow. (F-G) URX (F) and BAG (G) dendrite lengths quantified as a percentage of the distance from the cell body to the nose in the indicated genotypes. Each colored bar represents a single dendrite (n=50 per genotype); black bars indicate population averages. The shaded region marks wild-type mean ± 5 SD, and the percentage of dendrites in this range (“full-length”) is indicated below the plots. All quantification was performed on L4 larvae, except the far-right column which shows quantification for L1 larvae. Scale bars, 10 µm. p-values, Kruskal-Wallis with Dunn’s post hoc test adjusted for multiple comparisons.

MAGI-1 belongs to a family of proteins called <u>M</u>embrane-<u>A</u>ssociated <u>GU</u>anylate <u>K</u>inases (MAGUKs), which play key roles in organizing multiprotein complexes at cell-cell junctions (Zhu et al., 2016). In *C. elegans,* MAGI-1 localizes to epithelial junctions where it promotes the ordered arrangement of other junction components, and to synapses where it regulates postsynaptic receptor localization (Emtage et al., 2009; Lynch et al., 2012; Stetak and Hajnal, 2011; Stetak et al., 2009).

Consistent with these scaffolding functions, MAGI-1 contains a variety of protein-protein interacting domains characteristic of MAGUKs, including five PDZ domains, two WW domains, and a guanylate kinase domain that is catalytically inactive in MAGUKs and instead functions as a protein binding domain (Fig. 2A; Emtage et al., 2009). The mutant alleles we examined result in deletions that affect all known protein isoforms and give rise to frameshifts: *gk657* causes a small deletion-frameshift near the N-terminus, while *zh66* results in a larger 2.6kb deletion-frameshift in the first three PDZ domains (Fig. 2A) (*C. elegans* Deletion Mutant Consortium, 2012; Stetak et al., 2009). Importantly, these mutants are viable with grossly normal body morphology, suggesting that the URX and BAG dendrite defects are not secondary to broad defects in embryonic development.

*magi-1* mutants display more highly penetrant defects in URX than BAG (full-length dendrites in *magi-1(zh66)*: URX, 2%; BAG, 70%). Despite the modest penetrance of the BAG defects, affected BAG dendrites exhibit highly variable lengths (SD=36%, *magi-1(zh66*)), with some BAG neurons failing to extend any dendrite at all (n=4/50, *magi-1(zh66*)). By contrast, the URX phenotype is more consistent across individuals, with URX dendrites extending 66±13% and 65±16% of the distance from the cell body to the nose in *magi-1(gk657)* and *magi-1(zh66)* mutants, respectively. In this respect, *magi-1* phenotypically resembles *grdn-1,* which also affects URX more strongly than BAG, whereas *sax-7* affects BAG more strongly, although the significance of these differences remains unclear.

We decided to use the *zh66* allele throughout the remainder of this study because this large deletion is predicted to be a functional null (Stetak et al., 2009). URX and BAG dendrite defects are almost completely rescued by expression of MAGI-1a cDNA in the *magi-1(zh66)* mutant background (full-length dendrites: URX, 92%; BAG, 98%; Fig. 2F,G), confirming that dendrite morphogenesis defects in these mutants are due to disruption of the *magi-1* locus. We conclude that *magi-1* is required for URX and BAG dendrite extension.

### MAGI-1 functions during embryogenesis to promote dendrite extension

In principle, *magi-1* mutant phenotypes observed during late larval stages (L4) could result from defects in dendrite extension during embryonic development or from a failure to allometrically scale dendrites as the head grows ∼2-fold during larval development. To distinguish between these possibilities, we visualized URX and BAG in the embryo. We observed that URX and BAG dendrites are shortened in *magi-1* mutants during embryogenesis (Fig. 2D,E). Quantification of dendrite lengths relative to head size in newly hatched animals revealed a distribution of dendrite lengths that is similar to that observed in L4 animals (Fig. 2F,G, “*zh66* at L1” vs. “*magi-1(zh66)*”). We conclude that *magi-1* is required for the initial extension of URX and BAG dendrites during embryogenesis. The shortened dendrites then appear to scale allometrically with growth of the head during larval development.

### Mutations in *magi-1* preferentially affect URX and BAG dendrites

*magi-1* is widely expressed during embryonic development (Supp. Fig. 1), raising the possibility that mutations in this gene could cause broad defects in dendrite morphogenesis. To assess the specificity of *magi-1* dendrite extension defects, we examined *magi-1* mutants bearing cell-type-specific fluorescent markers for a panel of sensory neuron types that each extend a single, unbranched dendrite to the nose.

We first assessed a neuron called URY which – like URX and BAG – is not wrapped by glia and does not contact the external environment (Ward et al., 1975). We found that URY dendrite extension is not affected by mutations in *magi-1* (Fig. 3B).

**Figure 3.**
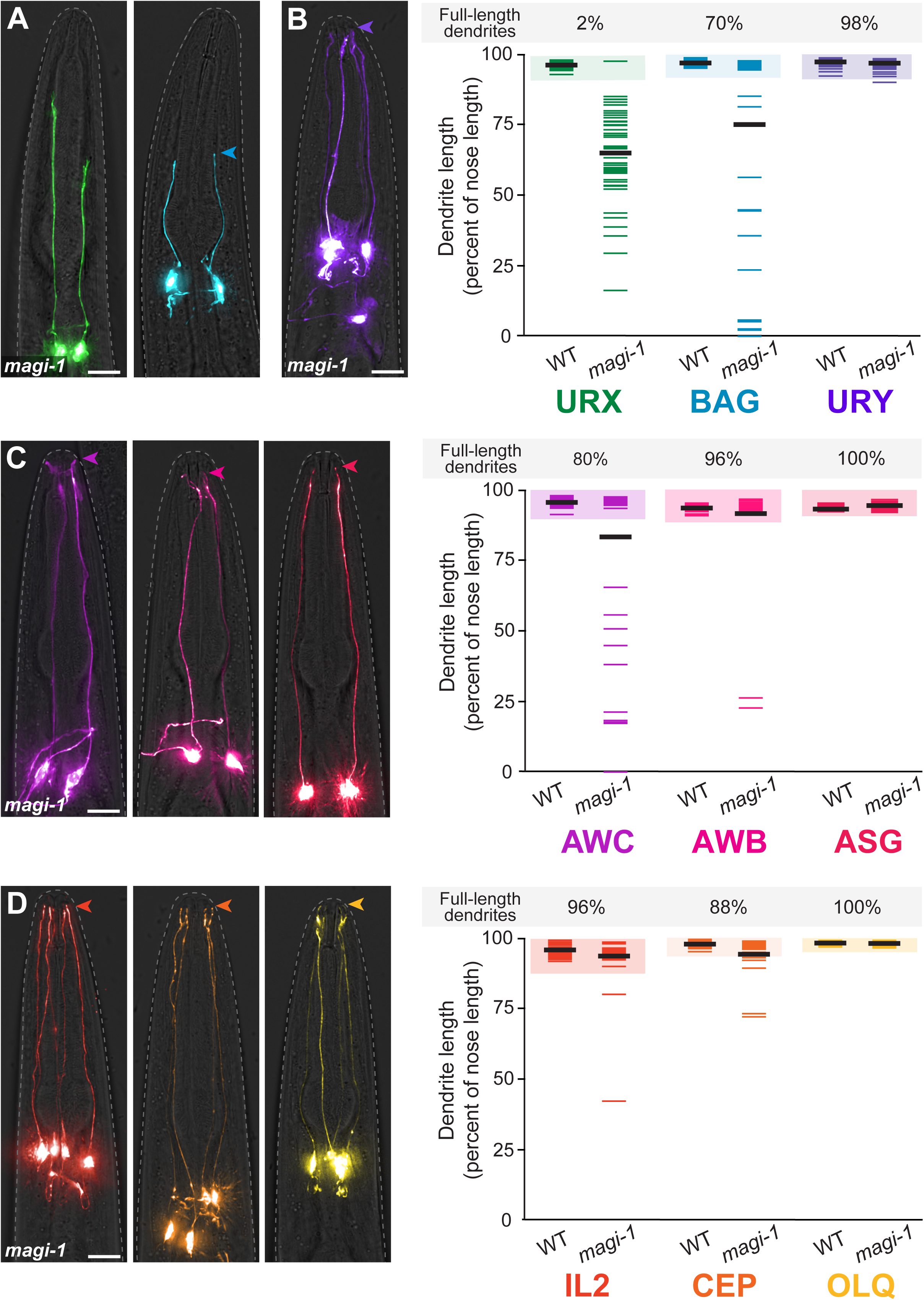
Mutations in *magi-1* preferentially affect URX and BAG dendrites. Dendrite lengths in *magi-1(zh66)* animals were measured for (A, B) neurons that do not protrude into the external environment, but form specialized attachments (A) to the ILso glia: URX (green, *flp-8*pro) and BAG (blue, *flp-17*pro) or (B) to other glia: URY (dark purple, *tol-1*pro) (Doroquez et al., 2014; Ward et al., 1975); (C, D) neurons that protrude through glial pores and form tight and adherens junctions with glia: (C) amphid neurons AWC (light purple, *odr-1*pro), AWB (pink, *str-1*pro), ASG (red, *ops-1*pro) and (D) neurons in other sense organs, IL2 (dark orange, *klp-6*pro), CEP (light orange, *dat-1*pro), OLQ (yellow, *ocr-4*pro). Arrowheads, dendrite endings. Scale bars, 10 µm. Quantification of dendrite lengths is shown at right, expressed as a percentage of the distance from cell body to nose. Colored bars represent individual dendrites (n=50 per genotype); black bars represent population averages. Shaded region represents wild-type mean ± 5 SD for each neuron type and the percentage of dendrites in this range (“full length dendrites”) is indicated above the plots. Images from (A) and quantification of URX and BAG dendrite lengths are reproduced from Fig. 2.

Next, we examined six sensory neurons that – unlike URX, BAG, and URY – each have a dendrite ending that forms a ring-shaped epithelial junction with a glial cell to demarcate a biochemically distinct apical region that protrudes into the external (or lumenal) environment (Low et al., 2019; Ward et al., 1975). These included three of the twelve neurons in the largest sense organ, the amphid (AWC, AWB, and ASG; Fig. 3C) and three neurons in other sense organs (IL2, CEP, OLQ; Fig. 3D).

Interestingly, we found that 20% of dendrites appeared shortened for the amphid AWC neuron, although the amphid AWB neuron – which has a similar structure – and the amphid ASG neuron were mostly unaffected (Fig. 3C). Dendrite extension defects in the other neuron types were also very rare (IL2, AWB, CEP) or absent (OLQ) (Fig. 3D). Morphological defects – shortening, as well as branching – have been observed in *grdn-1* mutants in individual amphid dendrites (AWA, AWB) (Nechipurenko et al., 2016), possibly due to disruption of the ring-shaped epithelial junctions with the amphid sheath glial cell that are required for dendrite extension. The defects upon loss of MAGI-1 may similarly reflect its known role in organizing epithelial junctions (Lynch et al., 2012; Stetak and Hajnal, 2011).

Thus, while *magi-1* plays a contributing role in dendrite extension for several neuron types that form ring-shaped epithelial junctions with glia, in general most neurons withstand its loss and show minor or no defects in dendrite extension. By contrast, URX and BAG dendrites do not form ring-shaped junctions with glia and are unusually sensitive to loss of MAGI-1. We reasoned that the difference in how they attach to glia may explain their differential sensitivity to loss of MAGI-1.

### MAGI-1 can act in glia to non-cell-autonomously promote URX dendrite extension

We next sought to determine in which cells *magi-1* acts to promote URX and BAG dendrite extension. We first considered the possibility that *magi-1* acts cell autonomously in URX and BAG. To test this hypothesis, we performed mosaic rescue analysis to interrogate the activity of MAGI-1 under the control of its own promoter. We focused on URX because the *magi-1* dendrite phenotypes are nearly 100% penetrant for this neuron.

Briefly, in *C. elegans,* genes of interest can be expressed through unstable extrachromosomal arrays that are stochastically lost during cell division. We generated *magi-1* animals carrying an array with *magi-1*pro:MAGI-1a cDNA, together with a URX-specific mCherry reporter to mark the presence of the array in URX. In this same line, we also labeled URX using a fluorescent GFP reporter that was stably integrated into the genome. We selected for mosaicism by picking animals in which only one of the two URX neurons was mCherry(+) (and therefore MAGI-1a(+)) and assessed dendrite phenotypes using GFP fluorescence. If MAGI-1 were required cell-autonomously in the neuron, we reasoned that all MAGI-1(+) URX neurons would have full-length dendrites and all MAGI-1(–) URX neurons would have dendrite extension defects (Fig. 4A, "Class I"). We found that only 2 of 36 mosaic animals exhibited a phenotype that matches this prediction (Fig. 4A). Instead, in 33 of 36 mosaic animals, both URX neurons extended full-length dendrites even though MAGI-1a cDNA was only present in one of the two neurons (Fig. 4A, “Class II”). We also saw one example in which the MAGI-1(+) neuron displayed dendrite defects and the MAGI-1(–) neuron was morphologically normal (Fig. 4A, “Class IV”). Therefore, MAGI-1 is not required cell autonomously in URX to promote dendrite development.

**Figure 4.**
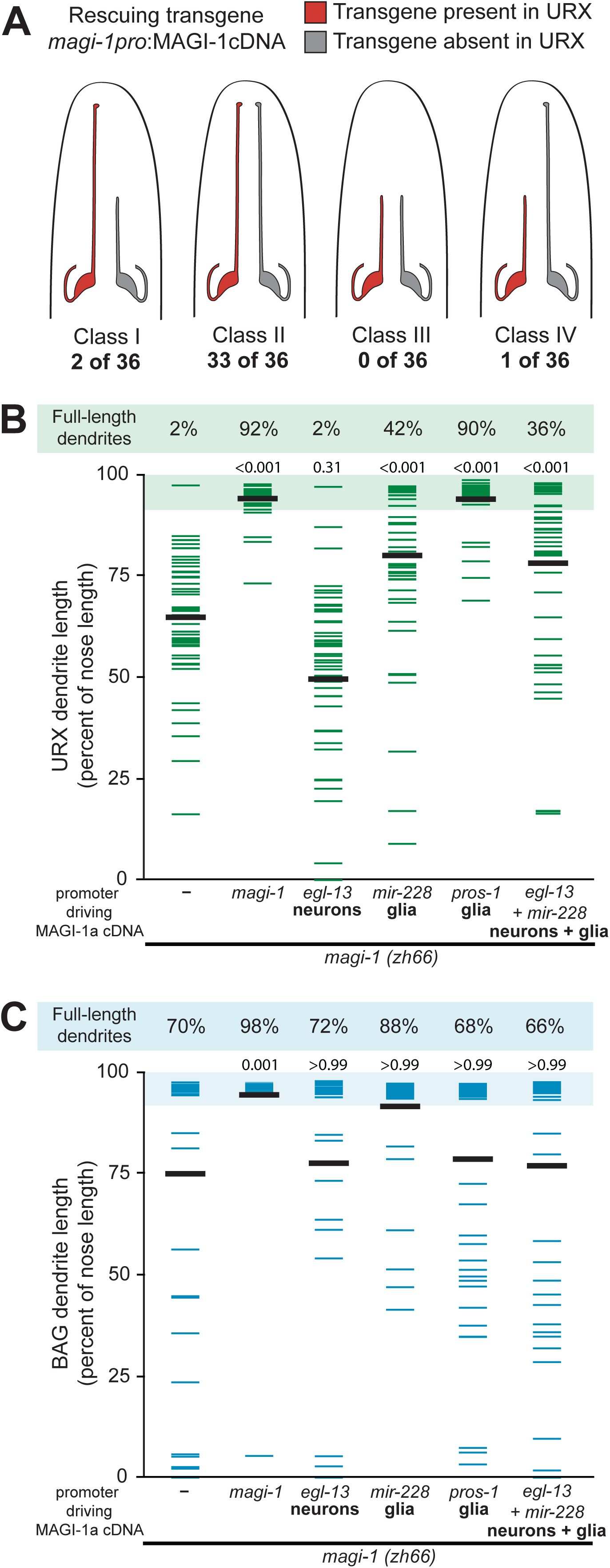
MAGI-1 can act in glia to non-cell-autonomously promote URX dendrite extension. (A) Schematic indicating results of a mosaic rescue experiment. Analysis was performed using *magi-1(zh66)* mutant animals bearing a stably integrated URX marker for assessing dendrite lengths (*flp-8*pro:GFP) and an unstable extrachromosomal array – stochastically lost during cell division – containing a rescuing MAGI-1 cDNA (*magi-1*pro:MAGI-1a) with a second URX marker for assessing the presence of the transgene in this neuron (*flp-8*pro:mCherry). To select for mosaicism, animals were selected in which only one of the two URX neurons was mCherry(+), indicating the presence of the rescuing cDNA, and the length of mCherry(+) (red) and mCherry(–) (gray) dendrites was assessed. The number of animals showing each of the four possible outcomes (Class I-IV) is indicated. (B-C) URX (B) and BAG (C) dendrite lengths, expressed as a percentage of the distance from the cell body to the nose, in *magi-1(zh66)* mutant animals bearing no transgene (–) or transgenes with a rescuing MAGI-1a cDNA under the control of the indicated promoters. Colored bars represent individual dendrites (n=50 per genotype); black bars represent population averages. Shaded region represents wild-type mean ± 5 SD and the percentage of dendrites in this range (“full length dendrites”) is indicated above the plots. *magi-1(zh66)* and +*magi-1*pro:MAGI-1 data are reproduced from Fig. 2. p-values, Kruskal-Wallis with Dunn’s post hoc test adusted for multiple comparisons.

We next performed cell-specific rescue experiments to further verify this result and extend our analysis to include BAG. We expressed the MAGI-1a cDNA under the control of a promoter (*egl-13pro*) that is expressed in a small number of neurons in the embryo, including URX and BAG shortly after these cells are born. We observed that neither URX nor BAG *magi-1* dendrite defects are rescued by this manipulation (Fig. 4B,C). We conclude that *magi-1* acts non-cell autonomously to promote both URX and BAG dendrite extension.

MAGUK proteins concentrate at sites of cell-cell contact (Zhu et al., 2016), and our previous work suggests that URX and BAG dendrite endings anchor to the nose during retrograde extension using attachments to glia (Cebul et al., 2020). Thus, we next considered the possibility that MAGI-1 might act in glia to non-cell-autonomously promote URX and BAG dendrite extension. We performed cell-specific rescue experiments as described above, this time using promoters that drive expression in glia. We found that expression of MAGI-1a under a broad glial promoter (*mir-228*pro, Fung et al., 2020; Pierce et al., 2008) produces moderate rescue of URX dendrite development defects (full-length dendrites: 42% vs. 2%) (Fig. 4B). We observed a similar trend for BAG (full-length dendrites: 88% vs. 70%; Fig. 4C), though the degree of rescue did not rise to statistical significance; because the penetrance of BAG dendrite extension defects in *magi-1(zh66)* animals is already low, it is more challenging to detect incomplete rescue. Importantly, simultaneous expression of MAGI-1a in glia and neurons (*mir-228*pro + *egl-13*pro) did not rescue dendrite defects any more than expression in glial cells alone (Fig. 4B,C), consistent with the idea that MAGI-1 is not required cell-autonomously in URX and BAG. Instead, our results suggest that MAGI-1 can act in glia to non-cell autonomously promote dendrite extension.

The incomplete rescue we observed using the *mir-228* promoter could reflect differences in the developmental timing of expression compared to endogenous *magi-1* or a requirement for *magi-1* in other cells. To further test if glial MAGI-1 is sufficient to rescue dendrite extension phenotypes, we examined the rescuing activity of additional promoters that drive expression in subsets of glia. Interestingly, URX dendrite defects were nearly fully rescued by expression of MAGI-1a under the *pros-1* promoter, which labels a subset of glia and their precursors during embryogenesis (Fig. 4B) (full-length dendrites: 90% vs. 2%). Curiously, however, this same promoter did not rescue BAG dendrite extension phenotypes at all (Fig. 4C) (full-length dendrites: 68% vs. 70%). Given that URX and BAG extend their dendrites concurrently, this may indicate that these two neurons associate with different glial cells during dendrite extension stages. Altogether, our results are consistent with the idea that MAGI-1 can act in glia to promote URX and BAG dendrite extension.

### MAGI-1 and SAX-7 directly bind each other, yet have partially independent functions in dendrite extension

MAGI-1 has previously been implicated as a binding partner of SAX-7 (Lynch et al., 2012). Briefly, MAGI-1 localizes to epithelial cell-cell contacts in the developing *C. elegans* embryo, and this localization partly depends on SAX-7 (Lynch et al., 2012; Stetak and Hajnal, 2011). Moreover, in a yeast two-hybrid assay, MAGI-1 PDZ domains interacted with the SAX-7 cytoplasmic tail in a manner partly dependent on the SAX-7 PB motif (Lynch et al., 2012). We therefore investigated physical and genetic interactions between MAGI-1 and SAX-7 in the context of URX and BAG dendrite extension.

To test for direct binding, we performed pull-down experiments using purified SAX-7 cytoplasmic tail (isoform A, aa 1054-1144) fused to glutathione *S*-transferase (GST) and lysates of HEK293T cells expressing full-length FLAG-tagged MAGI-1 (Fig. 5A) or purified bacterially-expressed HIS-tagged MAGI-1 (Fig. 5B). This revealed a physical association between the two proteins (Fig. 5A,B, GST-SAX-7 WT). Because MAGI-1 is a multi-PDZ domain-containing protein, we further tested whether the SAX-7 PB motif is required for binding. Indeed, we found that deletion of the SAX-7 PB motif almost completely abolishes binding of the two proteins (Fig. 5A,B, GST-SAX-7 ΔPB), consistent with previous reports using the yeast two-hybrid assay (Lynch et al., 2012). Taken together, these results indicate that SAX-7 and MAGI-1 directly bind each other via the SAX-7 PB motif (Fig. 5C).

**Figure 5.**
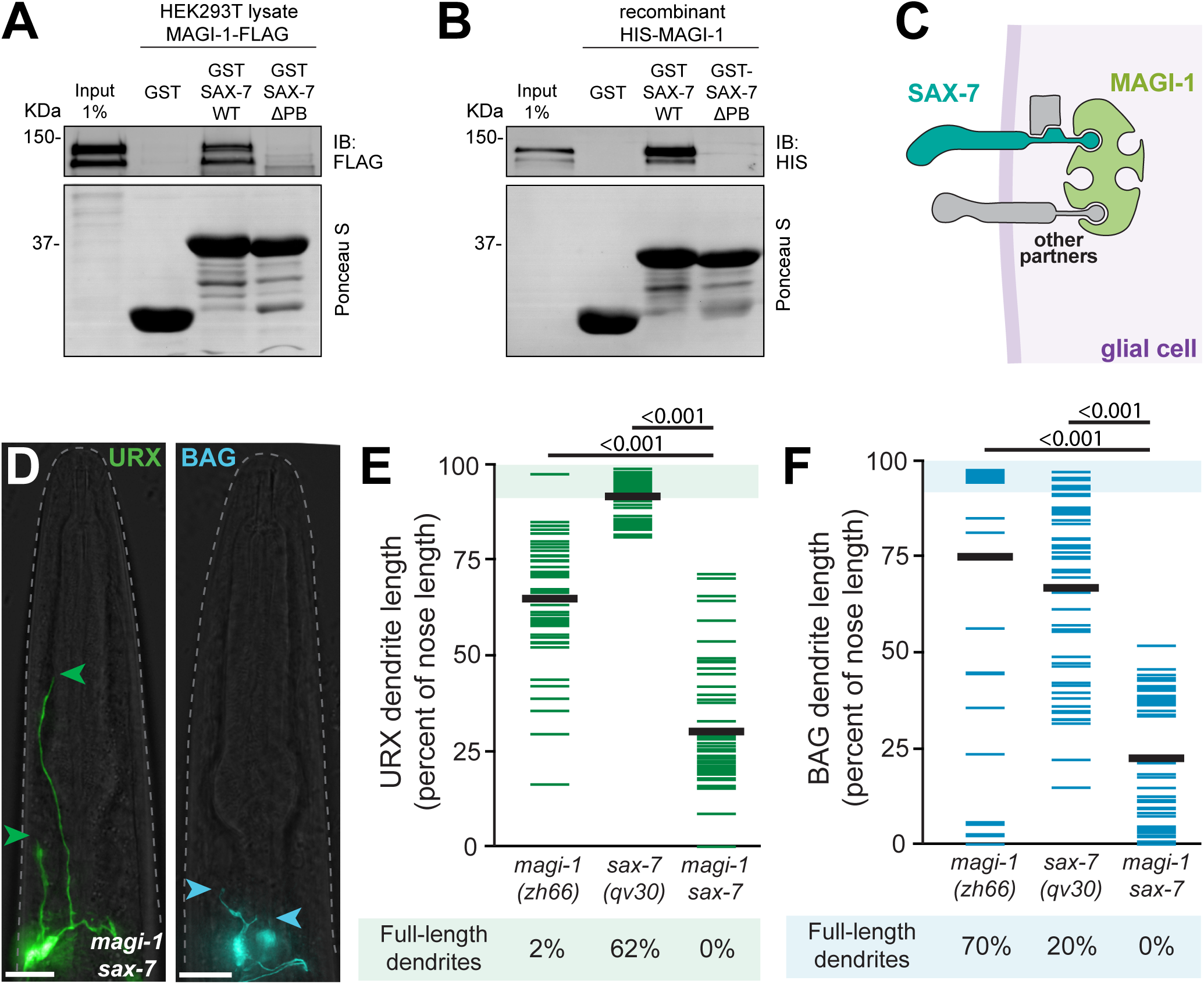
MAGI-1 and SAX-7 directly bind each other, yet have partially independent functions in dendrite extension. (A-B) Immunoblots from pull-down experiments. Glutathione *S*-transferase (GST) was fused to either the wild-type SAX-7 cytoplasmic tail (GST-SAX-7 WT) or the SAX-7 cytoplasmic tail lacking the PDZ-binding motif (GST-SAX-7ΔPB). These purified components were used in pull-down experiments with lysates of HEK293T cells expressing full-length FLAG-tagged MAGI-1 (A) or purified bacterially-expressed HIS-tagged MAGI-1 (B). Following pull-down, MAGI-1 was detected by immunoblot (IB) for the FLAG or HIS epitopes and the GST-tagged SAX-7 cytoplasmic tail was detected by Ponceau S staining for total protein. Each pull-down was performed at least twice with similar results. (C) Schematic indicating the physical interaction between SAX-7 (blue) and MAGI-1 (green). Other hypothesized interactors are indicated in gray. (D) Images of *magi-1(zh66) sax-7(qv30)* L4 larvae expressing cell-specific reporters for URX (green, *flp-8*pro) and BAG (blue, *flp-17*pro) neurons. Anterior, up. The dashed line outlines the head. Green and blue arrowheads, URX and BAG dendrite endings, respectively. Scale bars, 10 µm. (E-F) URX (E) and BAG (F) dendrite lengths quantified as a percentage of the distance from the cell body to the nose in the indicated genotypes. Each colored bar represents a single dendrite (n=50 per genotype); black bars indicate population averages. The shaded region marks wild-type mean ± 5 SD, and the percentage of dendrites in this range (“full-length”) is indicated below the plots. Wild-type and single mutant data are reproduced from Fig. 1 and 2. p-values, Kruskal-Wallis with Dunn’s post hoc test adjusted for multiple comparisons.

We next examined genetic interactions between *sax-7* and *magi-1 in vivo*. Using the previously reported amorphic alleles *sax-7(qv30)* and *magi-1(zh66)* (Desse et al., 2021; Stetak et al., 2009), we generated the *sax-7 magi-1* double mutant and examined URX and BAG dendrite lengths. In a simple model in which MAGI-1 function depends entirely on SAX-7 binding, we would predict that the phenotype of *sax-7 magi-1* double mutants would be no more severe than that of the single mutants. Instead, we found that dendrite extension defects are far more extreme in the double mutant as compared to either single mutant (Fig. 5D-F). Double mutant animals do not exhibit any full-length dendrites, and the average dendrite length is strongly reduced compared to *sax-7* and *magi-1* single mutants; for example, URX dendrites extend on average 30% of the distance from the cell body to the nose in double mutants, vs. 91% and 65% in *sax-7* and *magi-1* single mutants, respectively. These results suggest that SAX-7 and MAGI-1 are each able to promote dendrite extension in the absence of the other, likely through interactions with additional, partially redundant binding partners (Fig. 5C).

This idea is consistent with our previous observation that the SAX-7 AB, FB, and PB interaction motifs are required in combination, but not individually, to promote dendrite extension (Fig. 1). Similarly, previous work shows that multiple PDZ domains in MAGI-1 promote its junctional localization in the developing embryo in a manner that only partly depends on SAX-7 (Stetak and Hajnal, 2011). This suggests that SAX-7 and MAGI-1 may be part of a multi-protein adhesion complex that promotes dendrite extension, reminiscent of the large adhesive complexes at cell-cell junctions in epithelia. Strikingly, SAX-7, MAGI-1, and GRDN-1 have all been shown to associate with epithelial adherens junctions: *C. elegans* MAGI-1 and the GRDN-1 homolog in *Drosophila* physically interact with core components of adherens junctions (Houssin et al., 2015; Stetak et al., 2009), and both SAX-7 and MAGI-1 localize to epithelial junctions in *C. elegans* (Chen et al., 2001; Lynch et al., 2012). We thus wondered whether other components of epithelial adherens junctions also contribute to URX and BAG dendrite extension.

### The classical cadherin HMR-1 promotes URX and BAG dendrite extension

Classical cadherins are among the most prominent epithelial cell adhesion molecules. These highly conserved proteins are the core transmembrane component of epithelial adherens junctions and are present in all metazoans whose genomes have been analyzed (Nichols et al., 2012). The *C. elegans* genome encodes a single classical cadherin, called *hmr-1* (Costa et al., 1998; Hardin et al., 2013). HMR-1 and SAX-7 were previously shown to function redundantly during embryogenesis (Grana et al., 2010). Although strong loss-of-function alleles of *hmr-1* are lethal, previous work established tools to specifically deplete HMR-1 from cells of interest using the ZIF-1/ZF1 system (Armenti et al., 2014; Chihara and Nance, 2012). This approach allowed us to deplete HMR-1 from glia without causing widespread lethality, thus enabling examination of URX and BAG dendrite morphologies in these animals.

Briefly, we generated *hmr-1(zu389)* loss-of-function mutants bearing URX- or BAG-specific markers along with a functional, stably integrated *hmr-1* transgene tagged with a ZF1 degron sequence (Chihara and Nance, 2012; Costa et al., 1998; Von Stetina and Mango, 2015). The ZF1 tag targets proteins for rapid degradation in the presence of the E3 ubiquitin ligase substrate-recognition subunit ZIF-1 (Armenti et al., 2014). We then depleted HMR-1 from glia by expressing ZIF-1 under the control of a broad glial promoter, *mir-228*pro (Fung et al., 2020), which we previously used to rescue URX dendrite defects in *magi-1* mutants. These animals are viable, though some exhibit body morphology defects.

We found that HMR-1 glial knockdown animals exhibit dendrite extension defects in URX and BAG (Fig. 6A-D, HMR-1 KD in glia). URX is more strongly affected, with 54% of dendrites displaying morphological abnormalities, while the BAG phenotype is only weakly penetrant, with 12% of dendrites failing to extend fully. As a control, we found that this was not due to glial expression of ZIF-1 itself (Supp. Fig. 2C). We also examined IL2 neurons, whose dendrite endings are wrapped by the same glial cell that forms specialized attachments to the URX and BAG dendrite endings (Supp. Fig. 2A; Ward et al., 1975), and found that IL2 dendrites are unaffected by depletion of HMR-1 from glia (Supp. Fig. 2B). Notably, the strength of the URX and BAG phenotypes may underestimate the importance of HMR-1 in dendrite extension; although ZF1-mediated knockdown is reportedly efficient and rapid (within ∼30-45 minutes; Armenti et al., 2014), it remains possible that knockdown of HMR-1 is incomplete or delayed due to the kinetics of the *mir-228* promoter. Nonetheless, URX and BAG dendrites exhibit clear morphological defects upon glial knockdown of HMR-1. This result suggests that HMR-1 may mediate adhesion at the URX- and BAG-glia interfaces.

**Figure 6.**
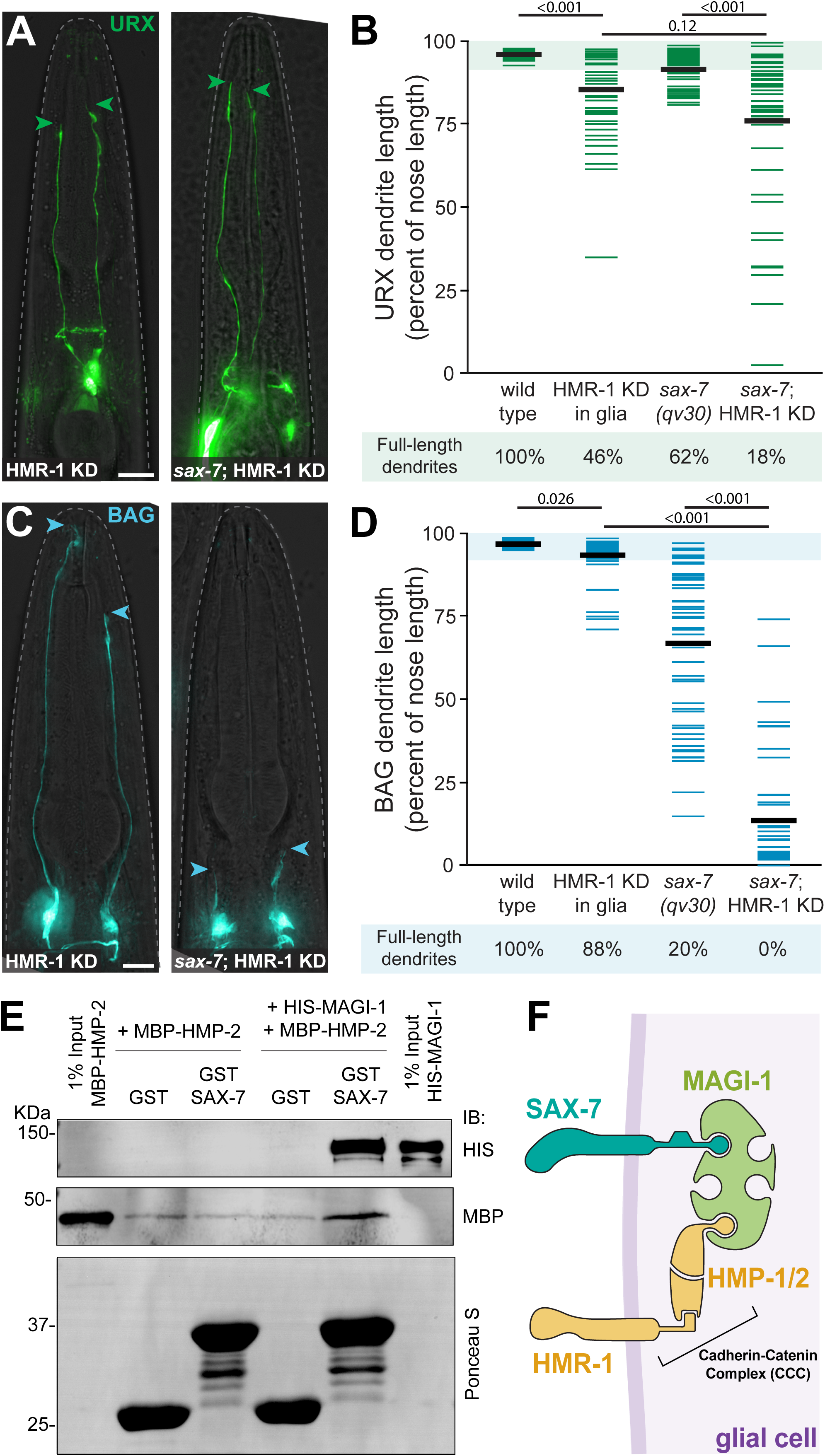
The classical cadherin HMR-1 promotes URX and BAG dendrite extension. (A,C) Images of URX (green, *flp-8*pro, A) and BAG (blue, *flp-17*pro, C) in wild-type (left) or *sax-7(qv30)* (right) animals with HMR-1 depleted from glia. Anterior, up. The dashed line outlines the head. Green and blue arrowheads, URX and BAG dendrite endings, respectively. Scale bars, 10 µm. Faint background fluorescent puncta are due to HMR-1-ZF1-GFP expression in non-glial cells. (B,D) URX (B) and BAG (D) dendrite lengths quantified as a percentage of the distance from the cell body to the nose in the indicated genotypes. “HMR-1 KD in glia” indicates knockdown of HMR-1 using the ZF1/ZIF-1 degron system for cell-specific degradation of HMR-1 using the pan-glial promoter *mir-228*pro. Each colored bar represents a single dendrite (n=50 per genotype); black bars indicate population averages. The shaded region marks wild-type mean ± 5 SD, and the percentage of dendrites in this range (“full-length”) is indicated below the plots. Wild-type and *sax-7(qv30)* data are reproduced from Fig. 1. Additional controls are shown in Supp. Fig. 2. (E) Immunoblot from pull-down experiments using purified recombinant components: the SAX-7 cytoplasmic tail fused to GST, HIS-tagged MAGI-1, and the HMP-2 PDZ-binding motif fused to MBP. Following pull-down, MAGI-1 and the HMP-2 motif were detected by immunoblot (IB) for the HIS or MBP epitopes respectively, and the GST-tagged SAX-7 cytoplasmic tail was detected by Ponceau S staining for total protein. Each pull-down was performed at least twice with similar results. (F) Schematic of MAGI-1 (green) bridging SAX-7 (blue) to the cadherin-catenin complex (gold) via HMP-2. p-values, Kruskal-Wallis with Dunn’s post hoc test adjusted for multiple comparisons.

Glia in *C. elegans* sense organs form canonical epithelial junctions (also called "apical junctions") with sensory neurons, other glia, and the skin (hypodermis) at the nose tip, such that each sense organ can be viewed as a miniature sensory epithelium continuous with the animal’s surface (Supp. Fig. 2A; Low et al., 2019; Ward et al., 1975). Each apical junction includes structures that are analogous to vertebrate adherens junctions and tight junctions (Labouesse, 2006; Pasti and Labouesse, 2014). Loss of HMR-1 could conceivably rupture these apical junctions, causing egregious defects in glial morphology that might have secondary effects on URX and BAG dendrite extension. We tested this possibility by visualizing ILso glia, together with URX, in HMR-1 knockdown animals. We found that even when URX neurons exhibit abnormal dendrite morphologies, ILso glia appear grossly normal (Supp. Fig. 2D). Because the ILso glia form both tight junctions and adherens junctions with the hypodermis, it is possible that the tight junctions can compensate for loss of HMR-1 at adherens junctions to stabilize glial morphology. In contrast, the URX- and BAG-glia contacts appear to lack tight junctions and, as such, likely lack this functional redundancy. In summary, the URX and BAG dendrite extension defects we observe are unlikely to be secondary effects of aberrant glial morphogenesis.

We conclude that HMR-1 can function in glia to non-cell autonomously promote the development of URX and BAG dendrites. Given the involvement of other factors (MAGI-1, SAX-7, GRDN-1) that are associated with epithelial adherens junctions, we propose that an adherens junction-like structure anchors the dendrite endings to glia at the nose in order to promote embryonic dendrite extension.

### HMR-1 functions redundantly with SAX-7 to promote URX and BAG dendrite extension

Previous work on early *C. elegans* embryonic development found that SAX-7 and HMR-1 function redundantly at sites of cell-cell contact to facilitate morphogenetic movements during gastrulation (Grana et al., 2010). We hypothesized that these two transmembrane cell adhesion molecules might also act redundantly in the context of URX and BAG dendrite extension.

To test this idea, we depleted HMR-1 from glia in *sax-7(qv30)* null animals and examined URX and BAG dendrite morphologies. Indeed, we found that knocking down HMR-1 in the absence of SAX-7 enhanced BAG dendrite extension defects as compared to *sax-7(qv30)* single mutants (Fig. 6A-D, *sax-7;* HMR-1 KD). For example, whereas 20% or 88% of BAG dendrites appear full-length in *sax-7* or HMR-1 knockdown animals, respectively, no full-length BAG dendrites were observed in *sax-7(qv30);* HMR-1 KD animals (Fig. 6D). We additionally noted a substantial enhancement in the expressivity of BAG dendrite phenotypes, with average dendrite lengths that were much shorter in double mutant animals (mean dendrite length: 67±24% in *sax-7*; 93±6% in HMR-1KD; 14±18% in *sax-7;* HMR-1KD) (Fig. 6D).

This effect was more subtle for URX. For example, 18% of URX dendrites still remained full-length in *sax-7(qv30);* HMR-1 KD animals (Fig. 6B). Dendrite lengths tended to be shorter in these animals compared to *sax-7(qv30*) (p<0.001) or HMR-1 KD alone (nominal p=0.02; adjusted p=0.13 after correction for multiple comparisons) although the effect was not as profound as for BAG (Fig. 6B,D). The observed differences between URX and BAG may reflect the presence of additional mechanisms that promote URX adhesion, or limitations in the timing of the HMR-1 depletion system relative to URX development.

Overall, these data are consistent with a model in which HMR-1 and SAX-7 function redundantly to promote URX and BAG dendrite extension.

### MAGI-1 can physically bridge SAX-7 to the cadherin-catenin complex

Classical cadherins interact with the cytoplasmic proteins HMP-1/α-catenin and HMP-2/β-catenin to form the cadherin-catenin complex (CCC) (Labouesse, 2006; Pasti and Labouesse, 2014), an essential feature of adherens junctions that links the transmembrane HMR-1/cadherin to the underlying actin cytoskeleton. Previous work on *C. elegans* MAGI-1 reported an interaction between MAGI-1 and HMP-2/β-catenin, mediated by the fifth PDZ domain of MAGI-1 and a PB motif at the C-terminus of HMP-2 (Stetak et al., 2009). Similar results have been reported for mammalian homologs (Dobrosotskaya and James, 2000). Given that the first three PDZ domains of MAGI-1 have been implicated in binding the cytoplasmic tail of SAX-7 (Lynch et al., 2012), we considered the possibility that MAGI-1 could form a ternary complex by simultaneously binding the SAX-7 and HMP-2/β-catenin PB motifs.

To test this hypothesis, we performed pull-down experiments using purified components: the SAX-7 cytoplasmic tail (isoform A aa 1054-1144) fused to GST, HIS-tagged MAGI-1, and the HMP-2 PB motif fused to MBP. In the absence of MAGI-1, pull-down of GST-SAX-7 did not enrich for MBP-HMP-2 compared to a GST-only control (Fig. 6E). By contrast, in the presence of HIS-MAGI-1, pull-down of GST-SAX-7 enriched for HIS-MAGI-1 as expected, as well as MBP-HMP-2 (Fig. 6E). This suggests that MAGI-1 can form a bridge that connects SAX-7 and HMP-2, likely using distinct PDZ domains to bind their PB motifs (Fig. 6F).

Altogether, our data indicate that MAGI-1 may promote robust URX- and BAG-glia adhesion by two potential mechanisms, which are not mutually exclusive. First, MAGI-1 may couple SAX-7 to the actin cytoskeleton via HMP-2 (Fig. 6F). This could fortify SAX-7-based attachments against mechanical breakage, similar to the role of actin coupling at cadherin-catenin complexes. Second, MAGI-1 may strengthen neuron-glia adhesion by clustering cell adhesion molecules, including SAX-7 and HMR-1, on the surface of the glial cell. By bridging SAX-7 and cadherin-catenin complexes, MAGI-1 scaffolding could lead to formation of a stronger adhesive network. Notably, this clustered adhesion model is consistent with the partial redundancy observed among individual adhesion components.

### The afadin homolog AFD-1 contributes to BAG dendrite extension

Finally, having identified a role for the epithelial cell adhesion molecule HMR-1/cadherin in URX and BAG dendrite extension, we tested whether other intracellular proteins associated with adherens junctions might also promote dendrite extension during embryogenesis. The scaffolding protein afadin localizes to epithelial adherens junctions in many species and facilitates junctional linkage to actin (Mandai et al., 1997; Mandai et al., 2013; Sawyer et al., 2009; Yonemura, 2011). Intriguingly, work in mammals and in *Drosophila* hints that afadins may be particularly important for epithelial integrity in dynamic tissues (Mandai et al., 2013; Manning et al., 2019; Sawyer et al., 2009). In *C. elegans*, AFD-1/Afadin has been shown to synergize with mutations that impair cadherin-catenin complexes, to co-localize with MAGI-1 in embryonic epithelia, and to interact with MAGI-1 in a yeast two-hybrid assay (Lynch et al., 2012; Serre et al., 2023). We therefore tested whether AFD-1 might also contribute to URX and BAG dendrite extension.

Because AFD-1 contains a PDZ domain (Fig. 7A), we first asked whether it binds the PB motif of the SAX-7 cytoplasmic tail. To this end, we isolated cDNA corresponding to the longest AFD-1 isoform (AFD-1a), expressed FLAG-AFD-1a in HEK293T cells, and performed pull-down assays using cell lysates and purified GST-SAX-7 cytoplasmic tail with or without its PB motif. We observed a modest interaction between AFD-1a and the SAX-7 tail that requires the PB motif (Fig. 7B, GST-SAX-7 vs. GST-SAX-7ΔPB). This suggests that AFD-1 may act as an alternative partner to MAGI-1 in SAX-7 binding (Fig. 7C).

**Figure 7.**
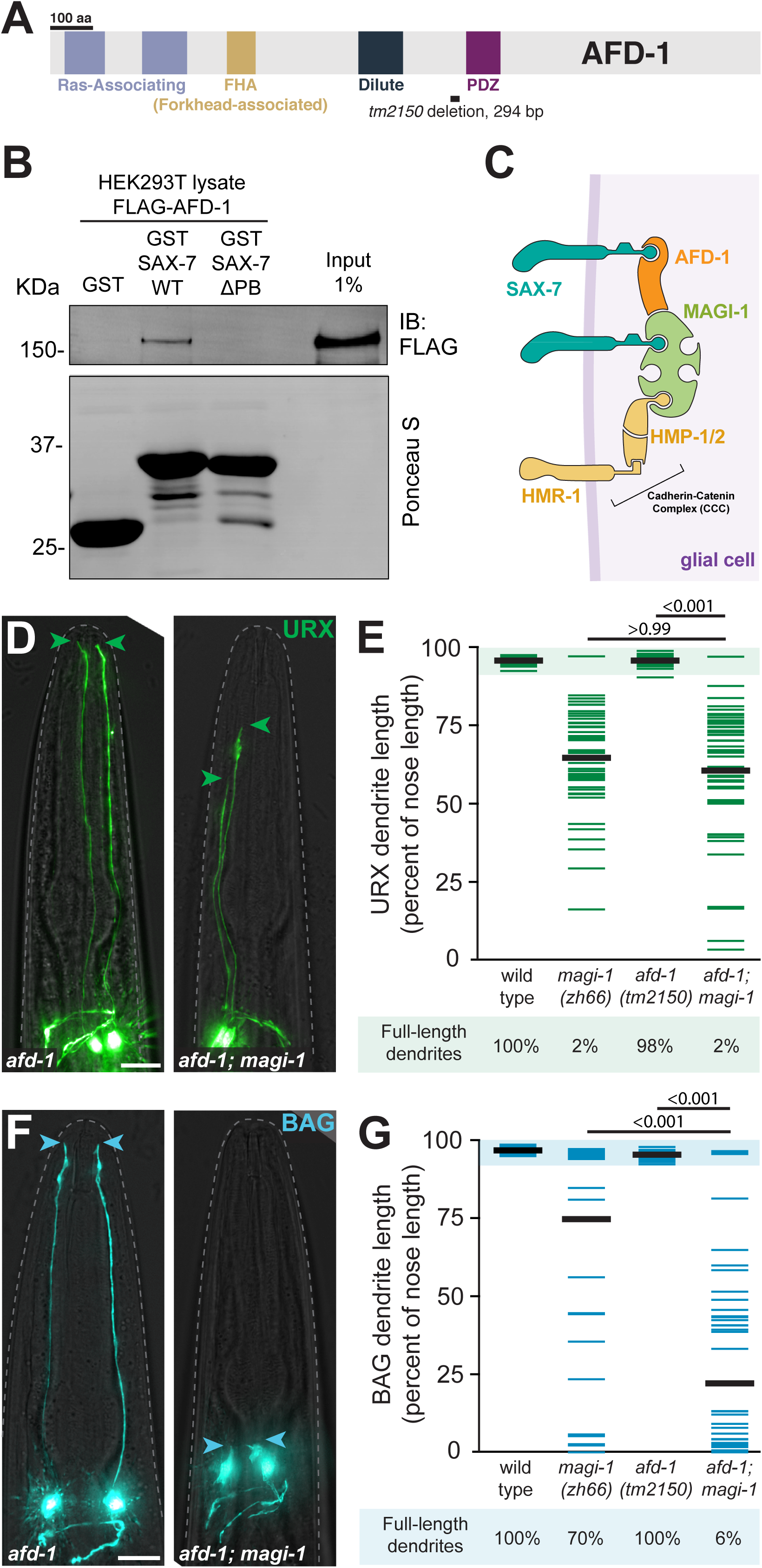
The afadin homolog AFD-1 contributes to BAG dendrite extension. (A) Diagram of AFD-1 indicating key protein domains and the allele used in this study. (B) Immunoblot showing pull-down experiments. GST was fused to either the wild-type SAX-7 cytoplasmic tail (GST-SAX-7 WT) or the SAX-7 cytoplasmic tail lacking the PDZ-binding motif (GST-SAX-7ΔPB). These purified components were used in pull-down experiments with lysates of HEK293T cells expressing full-length FLAG-tagged AFD-1. Following pull-down, AFD-1 was detected by immunoblot (IB) for the FLAG epitope and the GST-tagged SAX-7 cytoplasmic tail was detected by Ponceau S staining for total protein. Each pull-down was performed at least twice with similar results. (C) Schematic indicating two non-mutually-exclusive mechanisms for bridging SAX-7 to MAGI-1: AFD-1 (orange) connects SAX-7 (teal) to MAGI-1 (lime green) or SAX-7 directly binds MAGI-1, which in turn interacts with the cadherin-catenin complex via binding to HMP-2 (yellow). (D,F) Representative images of URX (green, *flp8*pro, D) and BAG (blue, *flp-17*pro, F) *afd-1(tm1250)* (left) or *afd-1(tm1250*)*; magi-1(zh66)* (right) mutant animals. Anterior, up. The dashed line outlines the head. Green arrowheads, URX dendrite endings. Blue arrowheads, BAG dendrite endings. Scale bars, 10 µm. (E,G) URX (E) and BAG (G) dendrite lengths quantified as a percentage of the distance from the cell body to the nose in the indicated genotypes. Each colored bar represents a single dendrite (n=50 per genotype); black bars indicate population averages. The shaded region marks wild-type mean ± 5SD, and the percentage of dendrites in this range (“full-length”) is indicated below the plots. *magi-1(zh66)* dendrite lengths are reproduced from Fig. 2. p-values, Kruskal-Wallis with Dunn’s post hoc test.

We next examined *afd-1(tm2150)* mutants, which contain a deletion-frameshift that is predicted to abolish expression of the C-terminal region that contains the PDZ domain (Fig. 7A; *C. elegans* Deletion Mutant Consortium, 2012). Although *afd-1(tm2150)* single mutants exhibit no deficits in dendrite development (Fig. 7D-G), we found that *afd-1(tm2150)* potently enhances BAG, but not URX, *magi-1* dendrite defects (Fig. 7D-G; BAG full-length dendrites: 6% vs. 70% in *magi-1(zh66)*). We observed that many double mutant animals have severely shortened BAG dendrites (average length: 22% vs. 75% in *magi-1(zh66)*). This result is consistent with a model in which AFD-1 and MAGI-1 act partially redundantly as alternative binding partners for SAX-7. It is not clear why URX is not further affected by *afd-1*, but this may reflect the strong effect of *magi-1* alone on URX dendrite extension. Indeed, nearly all URX dendrites exhibit morphological defects in *magi-1(zh66)* mutants alone. In summary, we find that AFD-1 binds the SAX-7 tail via its PB motif and acts redundantly with MAGI-1 to promote BAG dendrite extension (Fig. 7C).

## DISCUSSION

Proteins that mediate neuron-glia attachment are eagerly sought. Here, we find that MAGI-1 promotes URX and BAG dendrite extension during embryogenesis. Our results suggest that MAGI-1 functions in glia to anchor URX and BAG dendrite endings at the nose as dendrites develop. Mechanistically, we find that MAGI-1 directly binds both to the cytoplasmic tail of the cell adhesion molecule SAX-7 and to the cadherin-catenin complex protein HMP-2/β-catenin. Thus, MAGI-1 could bridge SAX-7 and HMR-1/cadherin together into a larger adhesion complex (Fig. 7C). Consistent with this idea, we find that both SAX-7 and HMR-1 also act in glia to non-cell-autonomously promote URX and BAG dendrite extension. This raises the possibility that URX and BAG dendrites initially attach to glia at the embryonic nose through a SAX-7-MAGI-1-HMR-1 adhesion complex.

Curiously, although our biochemical work suggests direct binding interactions between SAX-7, MAGI-1, and the cadherin-catenin complex, our genetic analysis reveals that each of these components can function independently of one another. For example, MAGI-1 remains active in the absence of SAX-7 (compare *sax-7* mutant to *magi-1 sax-7* mutant, Fig. 5). MAGI-1 therefore likely interacts with additional partners that contribute to dendrite extension, consistent with its identity as a multi-PDZ scaffolding protein. Similarly, SAX-7 remains at least partially active in the absence of MAGI-1 (compare *magi-1* mutant to *magi-1 sax-7* mutant, Fig. 5), indicating that its dendrite-anchoring activity does not require formation of a MAGI-1-based complex. Consistent with this notion, we find that the ankyrin-, FERM-, and PDZ-binding motifs in the SAX-7 tail function redundantly during dendrite development. Thus, SAX-7 likely promotes dendrite anchoring through interactions with additional binding partners via its ankyrin- and FERM-binding motifs. In addition, we find that the SAX-7 PDZ-binding motif binds to both MAGI-1 and an alternative partner, AFD-1/Afadin, that exhibits genetic redundancy with MAGI-1 in promoting BAG dendrite development (compare *magi-1* mutant to *afd-1; magi-1* mutant, Fig. 7). Together, the genetic redundancy and potentially multivalent interactions in this system suggest that it might be more useful to think of SAX-7, MAGI-1, AFD-1, and the cadherin-catenin complex as belonging to an ‘adhesion network’ at the glial surface, rather than a discrete adhesion complex.

The combined roles of MAGI-1, SAX-7, and HMR-1 in glia are highly reminiscent of their shared participation in adherens junctions in embryonic epithelia in *C. elegans,* and suggest that an adherens junction-like structure mediates neuron-glia attachment. During development of the *C. elegans* epidermis, partial loss-of-function mutations in the cadherin-catenin complex are strongly enhanced by depletion of MAGI-1, SAX-7, or AFD-1, leading to severe morphogenetic defects and embryonic lethality (Lynch et al., 2012). MAGI-1 and AFD-1 were shown to localize at or near adherens junctions in these cells, and this localization depends on SAX-7 (Lynch et al., 2012). Interestingly, depletion of MAGI-1 causes spatial disorganization of junction components (Lynch et al., 2012; Stetak and Hajnal, 2011), leading to the hypothesis that it acts as an organizer of other junction proteins. Homologs of MAGI-1 and AFD-1 have also been shown to play key roles at adherens junctions in *Drosophila* and in mammalian systems (Ikeda et al., 1999; Mandai et al., 1997; Mandai et al., 2013; Mizuhara et al., 2005; Sakakibara et al., 2020; Subauste et al., 2005). Taken together, these data suggest that URX and BAG dendrites anchor at the developing nose through a glial adherens junction-like structure.

The role of adherens junction proteins in URX and BAG dendrite development is surprising for two reasons. First, adherens junctions are most well-known for their role in attaching epithelial cells to one another (Niessen, 2007). In contrast, the mature URX and BAG dendrites are not part of an epithelium (Cebul et al., 2020; Ward et al., 1975), although we do not know the anatomical details of their initial attachments at the nose. Second, although *magi*-*1* and *sax-7* are broadly expressed, mutations in these genes preferentially affect URX and BAG dendrite extension. This is especially puzzling because other sensory dendrites – for example in the amphid – are embedded within sensory epithelia: they form bona fide tight junctions, exhibit apical-basal polarity, extend their dendritic endings into an external or lumenal apical environment, and require specific apical ECM proteins (e.g. DYF-7) for dendrite extension (Heiman and Shaham, 2009; Lillis et al., 2022; Low et al., 2019; Ward et al., 1975). Yet the effect of *magi-1* and *sax-7* mutations is milder or absent for these epithelial sensory dendrites, as well as for canonical epithelia like the epidermis (this study; Cebul et al., 2020). Thus, URX and BAG seem to have a greater susceptibility to mutations that, in epithelia, only mildly affect adherens junctions. An intriguing possibility is that URX and BAG dendrites attach to glia at the nose by adherens junction-like structures that are deployed outside the context of an epithelium, possibly akin to ‘fascia adherentiae’ or ‘puncta adherentiae’ that have been described ultrastructurally in other systems (Farquhar and Palade, 1963). These attachments might lack tight junctions, making them unusually susceptible to mutations that weaken the integrity of adherens junction complexes.

Interestingly, deletion of mouse Afadin/AFD-1 and mutations in human L1CAM/SAX-7 or CCDC88C/GRDN-1 all cause a shared phenotype: hydrocephalus, or enlargement of the ventricles in the brain (Adle-Biassette et al., 2013; Drielsma et al., 2012; Ekici et al., 2010; Ruggeri et al., 2018; Yamamoto et al., 2013). In the afadin conditional knockout mouse, hydrocephalus arises from loss of adherens junctions between the multiciliated ependymal cells that line the ventricle (Yamamoto et al., 2013), and defects in ependymal cell junctional complexes are increasingly thought to be a general mechanism of hydrocephalus (Rodríguez et al., 2012; Serra and Simard, 2023). Intriguingly, ependymal cells adhere to one another via adherens junctions, yet lack bona fide tight junctions after early embryonic development (Jiménez et al., 2014; Nelles and Hazrati, 2022; Serra and Simard, 2023). Thus, these cells – like URX and BAG – seem to be especially susceptible to mutations that weaken adherens junctions.

More broadly, it is intriguing to consider that adherens junction complexes without accompanying tight junctions may play a generalized role in dendrite-glia attachment. This idea is consistent with our understanding of synaptic contacts, as it has been suggested that epithelial adherens junctions are an evolutionary precursor of the synapse (Colman, 1999). Tight junctions are not necessary in this context, because there is no need to delineate internal and external surfaces within an inner neuropil. Interestingly, recent work on astrocyte-neuron interactions in visual cortex revealed that cadherin-catenin adhesion complexes between neurons and astrocytes regulate astrocyte morphogenesis, suggesting that modified adherens junctions may also be present at sites of neuron-glia contact in the mammalian brain (Tan et al., 2023). Thus, neurons and glia – which are developmentally derived from epithelia – may form attachments to one another by repurposing epithelial adherens junction machinery.

## METHODS

### Strains and plasmids

All *Caenorhabditis elegans* strains were maintained at 20°C on nematode growth media plates seeded with *E. coli* OP50 bacteria (Brenner, 1974). Mutant and transgenic lines were generated on the N2 background using standard techniques (Mello and Fire, 1995). Plasmids for bacterial production of GST-SAX-7, HIS-MAGI-1 and MBP-HMP-2 were generated by ligation-independent cloning (LIC) (Stols et al., 2002) of PCR-amplified fragments (SAX-7 aa 1054-1144; MAGI-1 aa 1-1054; HMP-2 aa 659-678) into pLIC-His, pLIC-GST, and pLIC-MBP (kindly provided by John Sondek, University of North Carolina, Chapel Hill), respectively. Plasmids for the expression of MAGI-1-FLAG (full-length C-terminally FLAG-tagged) and FLAG-AFD-1 (full-length N-terminally FLAG-tagged) in mammalian cells were constructed by insertion of full-length coding sequences between the NotI and EcoRI sites of p3XFLAG-CMV14 and p3XFLAG-CMV10, respectively, using Gibson assembly.

Detailed information on all strains generated in this study are presented in Supplementary Table 1; previously published strains are listed in Supplementary Table 2. Details on mutant alleles are provided in Supplementary Table 3, plasmids generated in this work are listed in Supplementary Table 4, and primers of general interest are included in Supplementary Table 5. Finally, amino acid sequences for SAX-7 rescue constructs are presented in Supplementary Table 6.

### Candidate screen for additional molecules required for URX and BAG dendrite extension

Mutations in known and predicted interactors of SAX-7 or GRDN-1, factors we previously identified through forward genetic screens for defects in dendrite morphology, were tested for URX and BAG dendrite extension defects. MAGI-1 was initially identified as a candidate because existing computational algorithms predicted that its PDZ domains interact with the SAX-7 PDZ-binding motif (Hui and Bader, 2010; Hui et al., 2013; Tonikian et al., 2008); in addition, Lynch and colleagues reported an interaction between MAGI-1 and SAX-7 via yeast two-hybrid (Lynch et al., 2012).

### Microscopy and image processing

Animals were prepared for imaging by mounting on a 2% agarose pad with a #1.5 coverslip. Larval animals were immobilized with sodium azide, young embryos (prior to twitching, ∼1.5-fold) were simply imaged live, and older embryos were arrested to prevent movement as described in Cebul, McLachlan, and Heiman, 2020. Early-stage embryos were acquired by dissecting adult hermaphrodites.

Image stacks were collected on a DeltaVision Core deconvolution imaging system (Applied Precision) with the InsightSSI light source; UApo 40×/1.35 NA oil immersion objective, PlanApo 60×/1.42 NA oil immersion objective or UPlanSApo 100×/1.40 NA oil immersion objective (Olympus); the standard DeltaVision live cell excitation and emission filter set; and a Photometrics CoolSnap HQ2 CCD camera (Roper Scientific). Image stacks were acquired and deconvolved with Softworx 5.5 (Applied Precision).

Representative images were generated as follows: Maximum intensity projections were created from contiguous optical sections in Fiji, then linearly adjusted for brightness in Adobe Photoshop. Multicolored images were generated by placing each channel in a separate false-colored screen layer in Photoshop. Adobe Illustrator was used to assemble final images.

### Dendrite length quantification

Dendrite lengths were measured at the L4 stage (except where otherwise noted) using the segmented line tool in Fiji. The starting point for tracing was defined as the junction between the dendrite and the cell body, and the final point was placed at the distal-most position along the dendrite. Dendrite lengths were then normalized by the distance between the cell body and the nose to account for size differences across animals. Dendrites were considered ‘full length” if they were within 5 standard deviations of the mean, as measured in L4 wild-type larvae (URX: >91.3%; BAG: >91.8%).

### Mosaic analysis of MAGI-1 activity

Animals of genotype *magi-1(zh66)* IV; *ynIs78* X (*flp-8*pro:GFP) carried an unstable extrachromosomal transgene array with *flp-8*pro:mCherry and a *magi-1*pro:MAGI-1a construct that almost completely rescues dendrite extension phenotypes. To enrich for mosaic animals, we selected animals in which one URX dendrite carried the array and one did not, scored using only the *flp-8*pro:mCherry marker on a fluorescence-equipped dissecting stereomicroscope. Each mosaic animal was then examined by GFP fluorescence, and dendrites with and without the array were scored as “full-length” or “short”.

### Protein expression and purification

HIS-tagged, GST-tagged, and MBP-tagged proteins were expressed in BL21(DE3) *Escherichia coli* transformed with the corresponding plasmids by overnight induction at 23°C with 1 mM isopropyl- β-d-1-thio-galactopyranoside (IPTG). For MBP-tagged proteins, bacterial growth media was supplemented with 0.2% (weight/vol) glucose. Bacteria pelleted from 1 liter of culture were resuspended in 25 ml of buffer (50 mM NaH_2_PO_4_, pH 7.4, 300 mM NaCl, 10 mM imidazole, 1% [vol/vol] Triton X-100 supplemented with protease inhibitor cocktail [leupeptin 1 µM, pepstatin 2.5 µM, aprotinin 0.2 µM, phenylmethylsulfonyl fluoride 1 mM]). After four pulses of sonication (20 second each with 1 minute rest on ice), lysates were centrifuged at 12,000 × *g* for 20 min at 4°C. The soluble fraction of the lysate was used for affinity purification on HisPur Cobalt resin (Thermo, Cat#89964), Glutathione Agarose resin (Thermo, Cat#16100) or Amylose Resin (New England Biolabs, E8021), depending of the fusion tag, and eluted with 50 mM Tris-HCl (pH 8), 100 mM NaCl supplemented with either 250 mM imidazole, or 30 mM reduced glutathione, or 20 mM amylose, respectively. Proteins were dialyzed overnight at 4°C against PBS. All protein samples were aliquoted and stored at −80°C.

### *In vitro* protein-protein binding assays

GST-SAX-7 proteins (30 µg of total protein) or a matching amount of GST were immobilized on glutathione-agarose beads (Thermo, Cat#16100) in PBS for 90 min at room temperature. Beads were washed twice with PBS, resuspended in 300 µl of binding buffer (50 mM Tris-HCl, pH 7.4, 100 mM NaCl, 0.4% [vol/vol] NP-40, 10 mM MgCl_2_, 5 mM EDTA, 2 mM DTT), and incubated 4 h at 4°C with constant tumbling in the presence of purified HIS-MAGI-1 (40 µg), MBP-HMP-2 (10 μg), or cleared lysates of HEK293T cells expressing FLAG-AFD-1 or MAGI-1-FLAG. For the preparation of HEK293T lysates, one-fourth of a 10-cm plate lysed in cold lysis buffer (20 mM HEPES, pH 7.2, 5 mM Mg(CH_3_COO)_2_, 125 mM K(CH_3_COO), 0.4% [vol/vol] Triton X-100, 1 mM DTT, 10 mM β-glycerophosphate, and 0.5 mM Na_3_VO_4_ supplemented with a protease inhibitor cocktail [Sigma S8830]) was used for each pull-down reaction. Each 10-cm plate was transfected with 9 μg of the indicated plasmid DNA construct using the calcium phosphate method. After the 4h incubation, beads from all binding reaction were washed four times with 1 ml of wash buffer (4.3 mM Na_2_HPO_4_, 1.4 mM KH_2_PO_4_, pH 7.4, 137 mM NaCl, 2.7 mM KCl, 0.1% [vol/vol] Tween-20, 10 mM MgCl_2_, 5 mM EDTA, 1 mM DTT), and beads-bound proteins were eluted with Laemmli sample buffer by incubation at 37°C for 10 min. Proteins were separated by SDS-PAGE and transferred to PVDF membranes, which were blocked with 5% (w/v) nonfat dry milk and sequentially incubated with primary and secondary antibodies diluted in 2.5% (w/v) nonfat dry milk with 0.05% (w/v) sodium azide. PVDF membranes were stained with Ponceau S and scanned before blocking. Each pull-down experiment was performed in at least two independent replicates with similar results. The primary antibodies used were the following (dilution factor in parenthesis): mouse anti-FLAG (Sigma F1804, 1:2,000), anti-HIS (Sigma H1029 1:2,500) or anti-MBP (New England Biolabs E8032, 1:5,000), goat anti-mouse IRDye 800 (Li-Cor Biosciences 926-32210, 1:10,000). Infrared imaging of immunoblots was performed using an Odyssey CLx Infrared Imaging System (LI-COR). Images were processed using ImageJ software (National Institutes of Health) or Image Studio software (LI-COR), and assembled for presentation using Photoshop and Illustrator software (Adobe).

### Statistical analysis

Data was initially recorded in Microsoft Excel or Apple Numbers. Statistical analysis was later performed using GraphPad Prism 9. A minimum of 46 dendrites (23 animals) were quantified for each genotype. All replicates are biological. Because all datasets failed normality tests and included multiple comparisons, the non-parametric Kruskal-Wallis test was used. Dunn’s post hoc test was employed to examine pairwise comparisons using the multiplicity-adjusted p-values option in Prism to adjust for multiple comparisons.

## ACKNOWLEDGEMENTS

We thank Jeremy Nance, Jeff Hardin, and Shai Shaham for sharing strains and reagents, and Ikue Mori for *sax-7* cDNA. Some strains were provided by the CGC, which is funded by NIH Office of Research Infrastructure Programs (P40 OD010440). We acknowledge the International *C. elegans* Gene Knockout Consortium for generating strains used in this study and the following funding sources: the National Science Foundation Graduate Research Fellowship Program (E.R.C.), NIH F31NS103371 (E.R.C.), NIH R01NS112343 (M.G.H.), NIH R01GM136132 (M.G-M.), and R01AG041870 (C.Y.B.).

