## Supplemental Data for "SAX-7/L1CAM acts with the adherens junction proteins MAGI-1, HMR-1/Cadherin, and AFD-1/Afadin to promote glial-mediated dendrite extension"

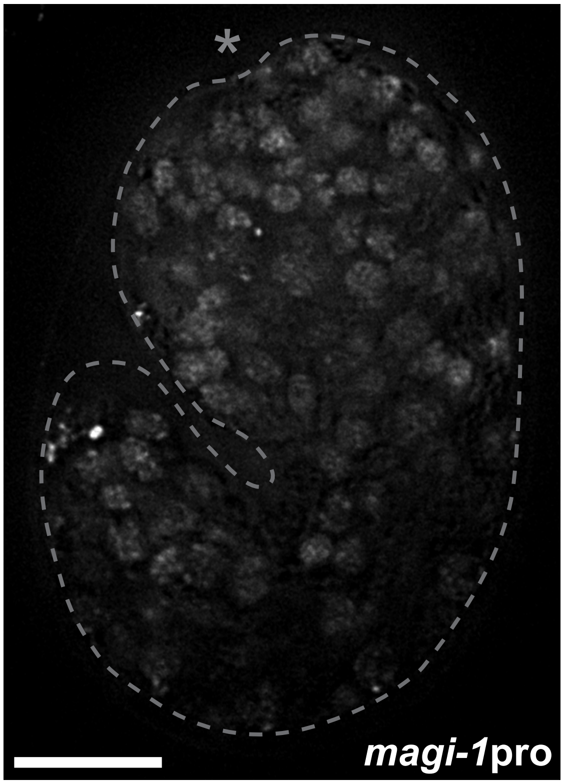


**Supplementary Figure 1. MAGI-1 is broadly expressed (Related to Fig. 3)**

A single optical section of a 1.5-fold embryo expressing nuclear histone-mCherry under control of the *magi-1* promoter, showing broad expression in the head. Dotted line outlines the embryo. Asterisk, nose. Anterior up, dorsal right. Scale bar, 10 µm.

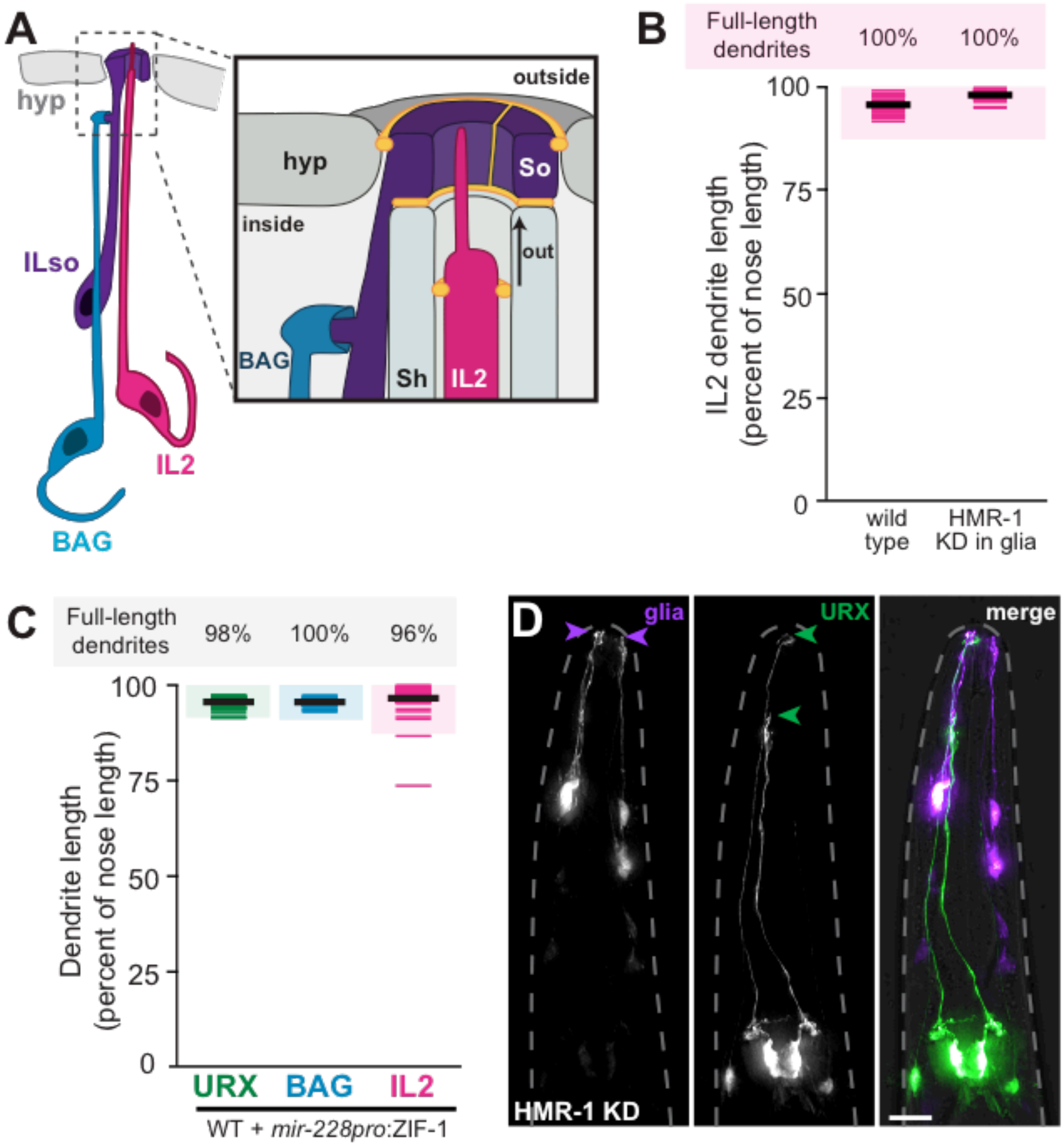


**Supplementary Figure 2. Glial knockdown of HMR-1 does not grossly affect ILso glial morphology or IL2 dendrite development. (Related to Fig. 6)**

(A) Schematic of the inner labial sense organ and its relationship to the BAG dendrite. The URX dendrite has a similar anatomy. The IL2 neuron (pink) has a dendrite that protrudes into the external environment through a pore formed by the IL sheath (Sh) and IL socket (So, purple) glia with tight and adherens junctions (yellow) between the dendrite and the glia and between the glia and skin (hyp). (B)

Quantification of IL2 dendrite lengths in wild-type or HMR-1 glial knockdown animals, as a percentage of the distance from the cell body to the nose. Wild-type data is reproduced from Fig. 3. (C) Controls for effects of glial ZIF-1 expression. URX, BAG, and IL2 dendrite lengths in animals bearing the same *mir-228pro*:ZIF-1 extrachromosomal array used for glial knockdown of HMR-1, but with wild-type *hmr-1* not subject to depletion. Dendrite lengths are quantified as a percentage of the distance from the cell body to the nose. (B-C) Each colored bar represents a single dendrite (n=50 per genotype); black bars indicate population averages. The shaded region marks wild-type mean ± 5 SD, and the percentage of dendrites in this range ("full-length") is indicated above the plots. (D) Glia (purple, left), URX (green, middle), or merge (right) in an animal with glial-specific knockdown of HMR-1. Glial endings remain at the nose as in wild-type animals (glial endings: purple arrowheads) even when URX dendrites are truncated (dendrite endings: green arrowheads). Scale bar, 10 μm.

**Supplementary Table 1. Strains generated in this study.**

| ID | Genotype | Figures |
|---|---|---|
| CHB2515 | *sax-7(qv30)*; *oyIs82*[*flp-17*pro:GFP + *unc-122*pro:dsRed] | 1, 5, 6 |
| CHB2508 | *sax-7(qv30)*; *ynIs78*[*flp-8*pro:GFP] | 1, 5, 6 |
| CHB3675 | *sax-7(qv30)*; *hmnEx2085*[*grdn-1*pro:SAX-7ΔFB + pRF4]; *ynIs78*[*flp-8*pro:GFP] | 1 |
| CHB3699 | *sax-7(qv30)*; *hmnEx2099*[*grdn-1*pro:SAX-7ΔAB + pRF4]; *ynIs78*[*flp-8*pro:GFP] | 1 |
| CHB3276 | *sax-7(qv30)*; *hmnEx1750*[*grdn-1*pro:SAX-7ΔPB + pRF4]; *ynIs78*[*flp-8*pro:GFP] | 1 |
| CHB3881 | *sax-7(qv30)*; *hmnEx2189*[*grdn-1*pro:SAX-7ΔFBΔABΔPB + pRF4]; *ynIs78*[*flp-8*pro:GFP] | 1 |
| CHB3821 | *sax-7(qv30)*; *hmnEx2168*[*grdn-1*pro:SAX-7minimal + pRF4]; *ynIs78*[*flp-8*pro:GFP] | 1 |
| CHB3730 | *sax-7(qv30)*; *hmnEx2085*[*grdn-1*pro:SAX-7ΔFB + pRF4]; *oyIs82*[*flp-17*pro:GFP + *unc-122*pro:dsRed] | 1 |
| CHB3820 | *sax-7(qv30)*; *hmnEx2099*[*grdn-1*pro:SAX-7ΔAB + pRF4]; *oyIs82*[*flp-17*pro:GFP + *unc-122*pro:dsRed] | 1 |
| CHB3542 | *sax-7(qv30)*; *hmnEx1750*[*grdn-1*pro:SAX-7ΔPB + pRF4]; *oyIs82*[*flp-17*pro:GFP + *unc-122*pro:dsRed] | 1 |
| CHB4062 | *sax-7(qv30)*; *hmnEx2189*[*grdn-1*pro:SAX-7ΔFBΔABΔPB + pRF4]; *oyIs82*[*flp-17*pro:GFP + *unc-122*pro:dsRed] | 1 |
| CHB3861 | *sax-7(qv30)*; *hmnEx2168*[*grdn-1*pro:SAX-7minimal + pRF4]; *oyIs82*[*flp-17*pro:GFP + *unc-122*pro:dsRed] | 1 |
| CHB2512 | *magi-1(zh66); ynIs78*[*flp-8*pro:GFP] | 2, 3, 4, 5, 7 |
| CHB2365 | *magi-1(zh66); oyIs82*[*flp-17*pro:GFP + *unc-122*pro:dsRed] | 2, 3, 4, 5, 7 |
| CHB2562 | *magi-1(gk657); ynIs78*[*flp-8*pro:GFP] | 2 |
| CHB2366 | *magi-1(gk657); oyIs82*[*flp-17*pro:GFP + *unc-122*pro:dsRed] | 2 |
| CHB2587 | *magi-1(zh66); ynIs78*[*flp-8*pro:GFP]; *hmnEx1435*[*magi-1*pro:MAGI-1a] | 2, 4 |
| CHB2551 | *magi-1(zh66); oyIs82*[*flp-17*pro:GFP + *unc-122*pro:dsRed]; *hmnEx1435*[*magi-1*pro:MAGI-1a] | 2, 4 |
| CHB3599 | *magi-1(zh66); hmnEx788*[*tol-1*pro:GFP + pRF4] | 3 |
| CHB3167 | *magi-1(zh66); oyIs44*[*odr-1*pro:dsRed] | 3 |
| CHB3233 | *magi-1(zh66); hmnEx1747*[*ops-1*pro:mCherry + *T02B11*pro:GFP + pRF4] | 3 |

| | | |
|---|---|---|
| CHB3168 | *magi-1(zh66); myIs13*[*klp-6*pro:GFP] | 3 |
| CHB3253 | *magi-1(zh66); kyIs104* [*str-1*pro:GFP] | 3 |
| CHB3204 | *magi-1(zh66); hmnEx782*[*dat-1*pro:GFP + *flp-8*pro:mCherry + pRF4] | 3 |
| CHB3254 | *magi-1(zh66); hmnEX641*[*ocr-4*pro:mCherry + pRF4] | 3 |
| CHB2623 | *magi-1(zh66); ynIs78*[*flp-8*pro:GFP]; *hmnEx1479*[*magi-1*pro:MAGI-1a + *flp-8*pro:mCherry] | 4 |
| CHB2687 | *magi-1(zh66); ynIs78*[*flp-8*pro:GFP]; *hmnEx1515*[*egl-13*pro:MAGI-1a + *flp-8*pro:mCherry + pRF4] | 4 |
| CHB2688 | *magi-1(zh66); oyIs82*[*flp-17*pro:GFP + *unc-122*pro:dsRed]; *hmnEx1515*[*egl-13*pro:MAGI-1a + *flp-8*pro:mCherry + pRF4] | 4 |
| CHB2737 | *magi-1(zh66); ynIs78*[*flp-8*pro:GFP]; *hmnEx1546*[*mir-228*pro:MAGI-1a + pRF4] | 4 |
| CHB3014 | *magi-1(zh66); oyIs82*[*flp-17*pro:GFP + *unc-122*pro:dsRed]; *hmnEx1546*[*mir-228*pro:MAGI-1a + pRF4] | 4 |
| CHB3575 | *magi-1(zh66); ynIs78*[*flp-8*pro:GFP]; *hmnEx2009*[*pros-1*pro:MAGI-1a + pRF4] | 4 |
| CHB3644 | *magi-1(zh66); oyIs82*[*flp-17*pro:GFP + *unc-122*pro:dsRed]; *hmnEx2009*[*pros-1*pro:MAGI-1a + pRF4] | 4 |
| CHB2833 | *magi-1(zh66); ynIs78*[*flp-8*pro:GFP]; *hmnEx1601*[*egl-13*pro:MAGI-1a + *mir-228*pro:MAGI-1a + pRF4] | 4 |
| CHB3020 | *magi-1(zh66); oyIs82*[*flp-17*pro:GFP + *unc-122*pro:dsRed]; *hmnEx1601*[*egl-13*pro:MAGI-1a + *mir-228*pro:MAGI-1a + pRF4] | 4 |
| CHB2873 | *sax-7(qv30) magi-1(zh66)*; *ynIs78*[*flp-8*pro:GFP] | 5 |
| CHB2884 | *sax-7(qv30) magi-1(zh66)*; *oyIs82*[*flp-17*pro:GFP + *unc-122*pro:dsRed] | 5 |
| CHB3159 | *hmr-1(zu389); xnIs375*[*hmr-1*-ZF1-GFP]; *hmnEx1646*[*mir-228*pro:ZIF-1-SL2-mCherry]; *ynIs78*[*flp-8*pro:GFP] | 6 |
| CHB3234 | *hmr-1(zu389); xnIs375*[*hmr-1*-ZF1-GFP]; *hmnEx1646*[*mir-228*pro:ZIF-1-SL2-mCherry]; *oyIs82*[*flp-17*pro:GFP + *unc-122*pro:dsRed] | 6 |
| CHB3822 | *hmr-1(zu389); xnIs375*[*hmr-1*-ZF1-GFP]; *hmnEx1646*[*mir-228*pro:ZIF-1-SL2-mCherry]; *sax-7(qv30); ynIs78*[*flp-8*pro:GFP] | 6 |
| CHB3698 | *hmr-1(zu389); xnIs375*[*hmr-1*-ZF1-GFP]; *hmnEx1646*[*mir-228*pro:ZIF-1-SL2-mCherry]; *sax-7(qv30); oyIs82*[*flp-17*pro:GFP + *unc-122*pro:dsRed] | 6 |
| CHB4042 | *afd-1(tm2150); ynIs78*[*flp-8*pro:GFP] | 7 |
| CHB2701 | *afd-1(tm2150); magi-1(zh66); ynIs78*[*flp-8*pro:GFP] | 7 |

| | | |
|---|---|---|
| CHB2513 | *afd-1(tm2150); oyIs82*[*flp-17*pro:GFP + *unc-122*pro:dsRed] | 7 |
| CHB2702 | *afd-1(tm2150); magi-1(zh66); oyIs82*[*flp-17*pro:GFP + *unc-122*pro:dsRed] | 7 |
| CHB3617 | *hmnEx2047*[*magi-1*pro:*his-24*-mCherry + *pros-1*pro:GFP + pRF4] | Supp. Fig. 1 |
| CHB3687 | *hmr-1(zu389); xnIs375*[*hmr-1*-ZF1-GFP]; *hmnEx1646*[*mir-228*pro:ZIF-1-SL2-mCherry]; *myIs14*[*klp-6*pro:GFP] | Supp. Fig. 2 |
| CHB3142 | *hmr-1(zu389); xnIs375*[*hmr-1*-ZF1-GFP]; *hmnEx1831*[*mir-228*pro:ZIF-1-SL2-mCherry + *grl-18*pro:YFP + *flp-8*pro:CFP] | Supp. Fig. 2 |
| CHB3146 | *hmnEx1646*[*mir-228*pro:ZIF-1-SL2-mCherry]; *ynIs78*[*flp-8*pro:GFP] | Supp. Fig. 2 |
| CHB3145 | *hmnEx1646*[*mir-228*pro:ZIF-1-SL2-mCherry]; *oyIs82*[*flp-17*pro:GFP + *unc-122*pro:dsRed] | Supp. Fig. 2 |
| CHB3679 | *hmnEx1646*[*mir-228*pro:ZIF-1-SL2-mCherry]; *myIs14*[*klp-6*pro:GFP] | Supp. Fig. 2 |

**Supplementary Table 2. Strains generated in previous studies.**

| ID | Genotype | Figure | Source/Reference |
|---|---|---|---|
| NY2078 | *ynIs78*[*flp-8*pro:GFP] | 1, 2, 3, 5, 6 | Kim and Li, 2004 |
| PY8503 | *oyIs82*[*flp-17*pro:GFP, *unc-122*pro:dsRed] | 1, 2, 3, 5, 6 | Astrid Cornils and Piali Sengupta |
| CHB1408 | *sax-7(hmn12)*; *ynIs78*[*flp-8*pro:GFP] | 1 | Cebul et al., 2020 |
| CHB1416 | *sax-7(hmn12)*; *oyIs82*[*flp-17*pro:GFP + *unc-122*pro:dsRed] | 1 | Cebul et al., 2020 |
| CHB1517 | *hmnEx788*[*tol-1*pro:GFP + pRF4] | 3 | Cebul et al., 2020 |
| PT2660 | *myIs13*[*klp-6*pro:GFP] | 3 | Schroeder et al., 2013 |
| PY1089 | *kyIs104*[*str-1*pro:GFP] | 3 | Troemel et al., 1997 |
| CHB1506 | *hmnEx782*[*dat-1*pro:GFP + *flp-8*pro:mCherry + pRF4] | 3 | Cebul et al., 2020 |
| CHB1304 | *hmnEx641*[*ocr-4*pro:mCherry + pRF4] | 3 | Cebul et al., 2020 |
| PY2417 | *oyIs44*[*odr-1*pro:dsRed] | Supp Fig 1 | Lanjuin et al., 2003 |
| CHB3098 | *hmnEx1747*[*ops-1*pro:mCherry + *T02B11*pro:GFP + pRF4] | Supp Fig 1 | Cebul et al., 2020 |
| PT2762 | *myIs14*[*klp-6*pro:GFP] | Supp Fig. 2 | Schroeder et al., 2013 |

**Supplementary Table 3. Mutant alleles used in this study.**

| Gene | Allele | Mutation Type | Reference |
|---|---|---|---|
| *sax-7* | *hmn12* | point mutation causing an early STOP | Cebul et al., 2020 |
| *sax-7* | *qv30* | CRISPR-generated deletion of sax-7 locus | Desse et al., 2021 |
| *magi-1* | *zh66* | 2.6kb deletion causing a frameshift | Stetak et al., 2009 |
| *magi-1* | *gk657* | 390 bp deletion causing a frameshift | *C. elegans* Deletion Mutant Consortium, 2012 |
| *hmr-1* | *zu389* | point mutation causing an early STOP | Costa et al., 1998 |
| *afd-1* | *tm2150* | 294 bp deletion causing a frameshift | *C. elegans* Deletion Mutant Consortium, 2012 |

**Supplementary Table 4. Plasmids generated in this study.**

| ID | Name |
|---|---|
| pEL172 | *egl-13*pro:MAGI-1a |
| pEL177 | *magi-1*pro:MAGI-1a |
| pEL278 | *magi-1*pro:his-24-mCherry |
| pEL205 | *mir-228*pro:MAGI-1a |
| pEL214 | *mir-228*pro:ZIF-1-SL2-mCherry |
| pEL238 | *grdn-1*pro:SAX-7ΔPB |
| pEL261 | *pros-1*pro:MAGI-1a |
| pEL263 | *pros-1*pro:GFP |
| pEL272 | *grdn-1*pro:SAX-7ΔFB |
| pEL273 | *grdn-1*pro:SAX-7ΔAB |
| pEL274 | *grdn-1*pro:SAX-7ΔFBΔABΔPB |
| pEL278 | *grdn-1*pro:SAX-7minimal |
| pLIC-GST | GST |
| pLIC-GST-SAX-7 | GST-SAX-7[1054-1144] |
| pLIC-GST-SAX-7ΔPB | GST-SAX-7[1054-1141] |
| pLIC-HIS-MAGI-1 | HIS-MAGI-1[1-1054] |
| p3XFLAG-MAGI-1 | MAGI-1[1-1054]-FLAG |
| pLIC-MBP-HMP-2 | MBP-HMP-2[659-678] |
| p3XFLAG-AFD-1 | FLAG-AFD-1[1-1565] |

**Supplementary Table 5. Primers of general interest.**

| Name | Sequence* |
|---|---|
| magi-1pro_fwd | CCTTCCTGCAGGTCGAACAGTTCACTCTTGC |
| magi-1pro_rev | CCTTGGCGCGCCCTGAATAAACTCATCGCCTCG |
| grdn-1pro_fwd | AGTATCAGCCTGCAGGCTTCGTAAATCTACAAAACATTTTCAACGTGC |
| grdn-1pro_rev | GAGATTCGGCGCGCCCCTTGATATTTTCGCTTGTTTTTTTTTCAGAAGG |
| egl-13pro_fwd | GATACCTGCAGGTGGGAGTTTGGTGCTTCC |
| egl-13pro_rev | GATAGGCGCGCCGTCTACGGCTGATGCTGG |
| pros-1pro_fwd | TAGCTCCTGCAGGGGTGATATCGAAAGTAACCAACG |
| pros-1pro_rev | TAGCTGGCGCGCCTGAGATTGATGACGTCACTAGC |
| mir-228pro_fwd | ATCACCTGCAGGGTGACGTCATACTCTTGC |
| mir-228pro_rev | TCAAGGCGCGCCAGTTTTTGGGAGGCGACG |

* Underlined nucleotides indicate restriction site.

**Supplementary Table 6. SAX-7 tail sequences.**

| Name | SAX-7 cytoplasmic tail sequence |
|---|---|
| WT | RQRGQNYPVSQREREQGREPILGKPDYKTDDDEKRSLTGSKAESETDSMAQYGDTDPGVFTEDGSFIGQYVPQKSLMPAERPEKGSTSTFV* |
| ∆FB | RQRGEQGREPILGKPDYKTDDDEKRSLTGSKAESETDSMAQYGDTDPGVFTEDGSFIGQYVPQKSLMPAERPEKGSTSTFV* |
| ∆AB | RQRGQNYPVSQREREQGREPILGKPDYKTDDDEKRSLTGSKAESETDSMAQYGDTDPGVFTEDGVPQKSLMPAERPEKGSTSTFV* |
| ∆PB | RQRGQNYPVSQREREQGREPILGKPDYKTDDDEKRSLTGSKAESETDSMAQYGDTDPGVFTEDGSFIGQYVPQKSLMPAERPE* |
| ∆FB∆AB∆PB | RQRGEQGREPILGKPDYKTDDDEKRSLTGSKAESETDSMAQYGDTDPGVFTEDGVPQKSLMPAERPEKGST* |
| minimal | RQRGEREQGREPILDELYK* |
